# Differential regulation of brain-specific molecular pathways is the reason for curcumin’s adult life-phase specific DAergic neuroprotection: Insights from ALSS Drosophila model of Parkinson’s disease

**DOI:** 10.1101/2025.01.17.633690

**Authors:** Abhik Das, Sarat Chandra Yenisetti

## Abstract

Curcumin (CU), a bioactive compound of turmeric, has been put forward as a “golden molecule” due to its anti-inflammatory, antioxidant, hepatoprotective, neuroprotective, and anti-cancer ability, as proven by research conducted over the decade and more. Our laboratory, developed an adult life stage specific (ALSS) *Drosophila* model of sporadic Parkinson’s disease (PD), and for first time demonstrated that dopaminergic (DAergic) neuroprotective efficacy of curcumin is limited to health phase viz. adult-young life stage and is absent during transition phase viz. adult senior life stages when PD set in. This observation suggests the limitation of curcumin as a therapeutic agent for late-onset disorders like PD. Further, our laboratory also demonstrated that despite curcumin’s ability to sequester oxidative stress during both the adult life stages, neuroprotection and brain dopamine replenishment is granted only in health stages but not in a vulnerable transition stage, which prompted to put forward the hypothesis that the molecular target(s) of CU, may be absent or inadequate in the transition stage of aging brain. With this insight, the current study was implemented to analyse the life stage-specific differential regulation of multiple molecular players of neuro-integral pathways in brain of ALSS *Drosophila* model of PD with curcumin intervention. It is discovered that curcumin-mediated health phase-specific neuroprotection underlies the correction of an altered expression of 1. *dFOXO, GADD45, Puc* of *Bsk-dFOXO* stress response pathway, 2. *Mfn2* of Mitochondrial dynamics 3. *CncC, GCLC, Prx 2540 -1,2, Jafrac1, Prx3* of Phase II antioxidant defense system pathway. Further, it is discovered that significant aging-associated naturally altered expression of certain molecular targets exists, that may contribute to the limitation of curcumin’s DAergic neuroprotective efficacy during the adult-transition stage. This knowledge will help in developing altered therapeutic strategies for PD as molecular targets of curcumin are conserved among fly, mice and human.

## 1. Introduction

The complicated interplay of gene and environment superimposed on slow and sustained neuronal dysfunction associated with aging [1,2] underlies cases of sporadic Parkinson’s disease (PD) in the population over the age of 60 [3]. The physiological symptoms are associated with motor and non-motor symptoms due to the death of dopaminergic (DAergic) neurons in the *SNpc* region of the human brain leading to the depletion of dopamine (DA) level [4]. Modern drug therapy of PD focuses on addressing the symptoms and consists mainly of dopamine (DA) supplementation through DA agonists like L-DOPA (Levodopa) or DA preservation through Catechol-o-methyltransferase and L-Monoamine Oxidase inhibitors [5]. As a part of combination therapy, L-DOPA-mediated DA supplementation is established as the gold standard treatment for PD and is the only therapeutic measure to counter late-onset symptoms of the disease [6]. However, L-DOPA-mediated therapy has been documented to be uncertain as it goes through “On-Off” phases, i.e., during the “On” phase patients respond to the therapy and during the “Off” phase patients do not respond to the therapy [7]. Further prolonged treatment with L-DOPA leads to the manifestation of “DA- resistant” motor, non-motor symptoms and neurotoxicity [8]. Therefore, symptomatic treatments are not enough to combat the onset and progression of PD. Hence, disease-modifying treatments such as nanoformulations, small molecules, immunotherapies, etc, are proposed as the new “cutting-edge tools” that may alter PD etiopathogenesis at the root level [6]. Multi-acting nutraceutical CU has gained potent interest in combating various chronic diseases including PD due to its genotropic property, i.e. ability to act on multiple molecular players [9].

In recent years, curcumin (CU) has also undergone human trials for PD. The efficacy of CU in human PD condition was also tested in a recent study. It was found that nano micelle CU treatment (80 mg/day for 9 months) on idiopathic PD patients (≥30 years of age) demonstrated significant improvement in a few aspects, overall, this did not culminate into statistically significant results for the whole trial [10]. The author concluded that although CU is a safe natural substance, this trial could not demonstrate its effectiveness in improving PD patients’ clinical symptoms and quality of life [10]. Similarly, CU’s limitation has been reflected in multiple chronic diseases in clinical trials [9]. Therefore, the inference is that the efficacy of CU in pre-clinical trials could not be translated to the human condition, and the current consensus is that future trials of any such nutraceuticals may directly be done in humans to cut short the discovery time and research expenditure [9]. However, it is important to note that *in-vitro* cell model studies might not recapitulate the *in-vivo* disease etiopathogenesis. Further, insights from the works in our laboratory suggest that when modeling a disease *in vivo*, variables such as life stages and their physiological implications on disease progression must be considered, so that the onset and progression of a disease match as near as possible to the human condition [11, 12]. Therefore, employing young animal models that do not match with appropriate life stages of disease onset in humans can be a possible reason for the failure to translate the results of preclinical studies to clinical setup.

With this insight, our lab focused on deciphering the neuroprotective efficacy of CU in an ALSS Drosophila model of sporadic PD developed in-house. The adult life of *Drosophila* is characterized by three stages, *viz.* health phase (HP) (No apparent mortality) corresponding to the adult young life stage, transition phase (TP) (10% mortality) corresponding to middle age of adult life and senescence phase (Steady decline in survival) corresponding to old age of adult life [13]. The aging of Drosophila is associated with alteration in genome-wide transcription profile by 23% [14]. Changes in genome-wide transcription profile during different life phases of Drosophila are similar to humans where significant aging-associated gene expression changes contribute to potent risk factors for an array of disorders like PD [15, 16]. Keeping the critical aspect that aging is a potential risk factor for late-onset NDD like PD, previously our laboratory developed the ALSS fly model and demonstrated that CU’s DAergic neuroprotective efficacy is HP-specific [12]. The same ALSS PD fly model is employed in the present study with an aim to understand the pathophysiology associated with PD before the occurrence of organismal death (employed neurotoxicant concentrations cause no mortality at the time points where phenotypes were scored) and to decipher the molecular basis of adult life phase specific CU’s DAergic neuroprotective efficacy.

To understand the implication of ALSS neuroprotective efficacy of CU in the PD model at neuronal and neurochemical levels, *in-situ* DAergic neurons and *ex-vivo* DA and metabolites were quantified with fluorescence microscopy and HPLC-ECD method. Our laboratory demonstrated that the onset of sporadic PD does not lead to degeneration of DAergic neuronal cell bodies but rather induces DAergic “neuronal dysfunction” during both HP and TP [under review]. “Neuronal dysfunction” is termed as the reduction of synthesis of tyrosine hydroxylase (TH), which is a rate-limiting enzyme for the synthesis of DA [17]. Further, diminished TH synthesis resulted in diminished DA level and the PD condition also prompted enhanced DA turnover, which may contribute to more summative oxidative stress [under review].

On the other hand, OS markers have been considered as the gold standard in the analyses of the neuroprotective efficacy of nutraceuticals since it is viewed as a contributing factor to NDD like PD [18–22]. In the PQ-induced ALSS (health and transition phase) PD fly model, Phom [23] quantified the oxidative stress (OS) in brain using OS markers viz., reactive oxygen species (ROS) level, protein carbonyl (PC) level, malondialdehyde (MDA) level, hydroperoxide (HPer) level, catalase (CAT) activity, superoxide dismutase (SOD) activity, glutathione s transferase (GST) activity, GSH level and neurotransmitter acetyl-cholinesterase (AChE) activity. It was demonstrated during both the adult life phases that levels of brain ROS, PC, HPer MDA and activity of CAT, SOD, GST were enhanced in induced PD condition. Further, exposure to PQ also showed downregulation of brain GSH level and AChE activity. However, CU intervention sequestered the upregulated ROS, PC, MDA, HPer levels, inhibited enhanced SOD, CAT, GST activity and rescued diminished GSH level, AChE activity [23]. It is important to note that though the brain OS is sequestered during both phases of adult life, CU fails to confer DAergic neuroprotection during TP, as is evident from its failure to rescue the brain DA level [12]. This insightful study illustrates that sequestration of oxidative stress may be necessary but is not sufficient to confer DAergic neuroprotection in PD. It is argued that apart from oxidative stress, other molecular players/pathways may have synergic effects that could be responsible for the observed DAergic neurodegeneration/neuroprotection [12]. Further, this evidence emphasizes the importance of employing life-stage matched animal models for late-onset NDD such as PD and stresses the limitation of OS and inflammation markers perse-based studies to determine DAergic neuroprotective efficacy of nutraceuticals/therapeutic agents/small molecules.

Herbicide paraquat (PQ) is a multi-hit environmental neurotoxicant whose underlying mechanism of dopaminergic (DAergic) neurodegeneration may underlie ROS generation, deficiency in antioxidant enzyme levels, neuroinflammation, mitochondrial dysfunction, and ER stress, leading to a cascade of molecular crosstalks that result in the initiation of apoptosis [24]. Therefore, to understand the mechanism of DAergic neuroprotective efficacy of CU, it is necessary to investigate the genetic/molecular players and their possible age-mediated differential regulation. It has been demonstrated that multiple molecular pathways are involved in the DAergic neuronal integrity. *JNK/BSK* signalling pathway and its downstream targets through *FOXO/dFOXO* axis is known for adaptive stress response and when regulated rightly can promote life span extension, neuroprotection [25–29]. IIS-*mTOR/dTOR* pathway is known for actively controlling anabolic processes and metabolism, impairment of such metabolic process known as “metabolic consequences” is a common phenomenon in onset and progress of PD [30–32]. The adaptive stress response controlled by *JNK/Bsk* signalling pathway and anabolic process controlled by IIS-*mTOR/dTOR* pathway influences mitochondrial dynamics which may also contribute to progression of NDD like PD [33–35]. Further, phase II ADS producing/utilizing GSH for antioxidant response [36, 37] and metal homeostasis pathway which prevents free metal accumulation in neuron [38, 39] are potent biological pathways that are involved in neuroprotection/neurodegeneration. Hence understanding the possible differential modulation of molecular players involved in these signalling networks may throw an opportunity to understand the ALSS neuroprotective efficacy of CU, understanding of which will be of great importance and support to develop novel treatment methods/regimen and alter existing therapeutic strategies

## 2. Materials and Methods

### 2.1. Modelling ALSS-PD in *Drosophila* and therapeutic intervention

Generation of the PQ-mediated early onset, late-onset sporadic PD model of *Drosophila* and therapeutic intervention with CU co-feeding is described in Phom et al [12]. The study of Phom et al [12] highlights the ALSS neuroprotective efficacy of CU. The current study uses the same model, where detailed description and results relating to selection of neurotoxicant and CU concentrations were provided [12].

#### 2.1.1. *Drosophila* stock and husbandry

Oregon K (OK) flies used in this study were obtained from the National *Drosophila* Stock Centre, University of Mysore, Mysore, Karnataka, India. Flies were maintained in food media containing (Sucrose, Agar-Agar, Yeast, and Propionic acid) [12, 40], under standard laboratory conditions of 22±2°C temperature with humidity of 60%, and 12:12 hrs light and dark condition.

#### 2.1.2. Collection and aging of adult male files

For the collection of adult male flies, the parental generation was transferred to a fresh media vial for laying eggs for 4 days and then removed. After 11-12 days the eclosed flies were lightly anaesthetised to separate males and females. Male flies were collected in food media vials (25 male flies in each vial) and aging was done according to the necessity of the experiment while switching them into fresh media vials every 4^th^ day. Flies belonging 4-5 days and 50-55 days representing HP and TP of adult life span respectively were used to model early and late-onset forms of PD as described in Phom et al [12].

#### 2.1.3. Chemicals for feeding and exposure

Sucrose (Cat. No. 1947139) procured from Sisco Research Laboratory (SRL, Mumbai, India), Type I Agar Agar (Cat. No. GRM666) procured from HiMedia (Thane, India), Propionic acid (Cat. No. 8006050-500-1730) procured from MERCK (Rahway, USA) and market-available sugar tolerant dry yeast (Angel, instant dry yeast) were used for food media preparation. For the exposure regimen, Methyl viologen dichloride hydrate /Paraquat (PQ) (Cat. No. 856177), CU (Cat. No.C1386) and Dimethyl Sulphoxide (DMSO) (Cat. No. D8418) were purchased from Sigma Aldrich (St. Louis, MO, USA). Feeding on Whatman filter paper no.1 disc in a 30x100 mm glass vial was preferred as the exposure methodology for the experiment.

#### 2.1.4. PQ treatment and CU co-treatment protocol

The necessary amount of PQ was dissolved in 5% sucrose solution to prepare 10 mM of PQ solution. A primary stock of 200 mg CU in 1 ml of DMSO was prepared (543 mM). Co-feeding of CU with PQ was achieved by dissolving 1.38 µL and 2.76 µL of CU stock in 1.5 ml of 10 mM PQ to obtain 500 µM and 1 mM of CU concentrations respectively. Also, exposure to CU *per se* at the concentrations (500 µM and 1 mM) was achieved by dissolving 1.38 µL and 2.76 µL of CU stock in a 5% sucrose solution only. Flies of the control group were fed 275 µL of 5% sucrose on filter discs while flies of experimental groups were fed 275 µL of 10 mM PQ (induced PD group), CU_500µM/1mM_ + 10 mM PQ (CU co-treatment group) and CU_500µM/1mM_ + 5% sucrose (CU *per se* group) respectively on filter discs. With 25 flies per vial, a minimum of 50-100 flies were exposed per group for 24 hrs [12].

### 2.2. Extraction of fly brain RNA and performing quantitative real time PCR (qRT-PCR) to quantify gene expression profiles

The methodologies pertaining to gene expression analysis utilizing qRT-PCR is partially described in Das et al [41].

#### 2.2.1. Chemicals and materials for gene expression analysis

RNaseZAP (Ambion, Waltham, USA, Cat: AM9780), DEPC treated water (HiMedia, Thane, India, Cat: ML024), TRIzol reagent (Invitrogen, Waltham, USA, Cat:15596026), Chloroform (Sigma-Aldrich, St. Louis, USA, Cat: c2432), Isopropyl alcohol (HiMedia, Thane, India, Cat: MB063-1l), DEPC treated water (HiMedia, Thane, India, Cat: ML024), Ethanol (MERK, Rahway, USA, Cat: 1.00983.0511), DNase (Invitrogen, Waltham, USA, Cat:18068-015), Oligo(dT) (Invitrogen, Waltham, USA, Cat: 18418012), dNTP mix (Qiagen, Hilden, Germany Cat: 14505289), SuperscriptTM II reverse transcriptase (Invitrogen, Waltham, USA, Cat: 18064014), PowerUp™ SYBR® Green Master Mix (Applied Biosystems, Waltham, USA, Cat: A25742), 96 reactions well plate (Biorad, Hercules, USA, Cat: MLL-9601).

#### 2.2.2. Total mRNA isolation from fly head

In brief, post-exposure each group of flies were frozen and heads were decapitated. 50 heads per fly group were used to extract total RNA extraction using Guanidium thiocyanate phenol chloroform extraction or the TRIzol method. Tissues were homogenized with 1 mL of TRIzol and centrifuged (At 12g for 15 mins at 2°C) to discard the tissue debris. Supernatant was collected and incubated at room temperature for 5 mins. Phase separation by chloroform (200 µL was added and vigorously shaken followed by incubation at room temperature for 2-3 mins) was opted to be the best way to isolate aqueous solution containing RNA from the homogenate. The tubes are centrifuged at 12g for 15 minutes at 2°C. Colourless upper aqueous phase was collected and RNA was eluted and precipitated out from the aqueous solution by adding chilled Isopropyl alcohol (500 µL was added and incubated at room temperature for 10 minutes followed by centrifugation at 13g for 15 min at 2°C). The pellet of RNA was washed with 1 mL of 75% ethanol to remove salt content from TRIzol reagent. Pellet was air dried and resuspended in 30-35 µL DEPC treated water (Incubated for 10 minutes at 55° to 60 °C to dissolve the pellet). RNA was quantified and stored in -80°C freezer till further use. Precaution taken to avoid RNase contamination, so every instrument and working space was cleaned with RNase ZAP prior to the experiment.

#### 2.2.3. First strand cDNA synthesis

RNA was quantified with pedestal method using Nanodrop (Thermo-Fisher scientific). 1 ug of RNA from each group is to be synthesized into single stranded cDNA. Required volume of RNA which contains 1 ug of the biomolecule was taken and treated with DNase+DNase buffer (1 µL each) to avoid DNA contamination. Volume makeup for each sample was done by adding DEPC water to obtain a uniform volume of 10 µL. Sample was incubated 15 mins at room temperature. After which 1 µL of EDTA was added followed by heating at 65°C for 10 minutes to deactivate the DNase. Tubes were puff centrifuged and Oligo DT and dNTP mix (1 µL each) were added followed by heating at 65°C for 5 minutes to promote annealing. Tubes were chilled immediately and then puff centrifuged. First strand buffer (4 µL) and 0.1 M DTT (2 µL) were added followed by heating at 42°C for 2 minutes to prepare the RNA template for first strand cDNA synthesis. 1 µL of Superscript II reverse transcriptase was added and incubated at 42°C for 50 minutes. The reaction was deactivated by incubating the tubes at 70°C for 15 minutes. Tubes were puff centrifuged again and now the first strand cDNA is ready which is stored in -20°C freezer till further use.

#### 2.2.4. Gene expression analysis

The Real time PCR reaction was carried out in Applied Biosystem (ABI) Step-One Plus thermal cycler (Thermo fisher scientific), using Power Up SYBR green master mix. cDNA was diluted with DEPC water with a factor 5 for the reaction. Reaction volume was considered 10 µl which contains 1 µl of diluted cDNA, 8.6 µl of SYBR green and 0.2 µl of forward and reverse primer specific to the internal control *RP49* and target gene *dFOXO*. Reaction plate was loaded accordingly as described in Das *et al*, 2021. The thermal cycler protocol for the qRT-PCR is as follows Amplification (Amp.) protocol: Holding stage @ 95°C C for 10 mins Amp. Cycle: (95°C for 15 secs followed by 60°C for 1 min) X 40 Melting Curve analysis: Quickly ramped up to 94°C followed by cooling at 60°C

The C_T_ values were obtained and the relative fold change to the Control sample was measured by the 2^-ΔΔCt^ method as described by [42].

For the experiment, in different treatment groups, utilizing the afore mentioned methodologies differential gene expression was analyzed for the molecular players belonging to *JNK* signaling pathway, *IIS-mTOR* signaling pathway, mitochondrial dynamics, Phase II antioxidant defense response and metal homeostasis (**Table 1**).

**Table 1:**
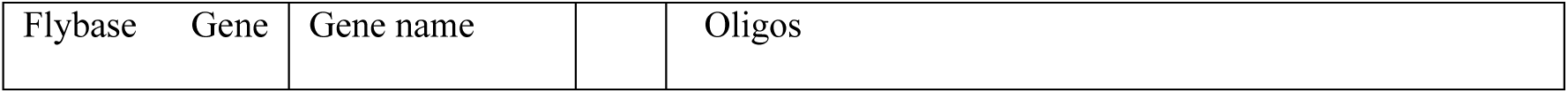
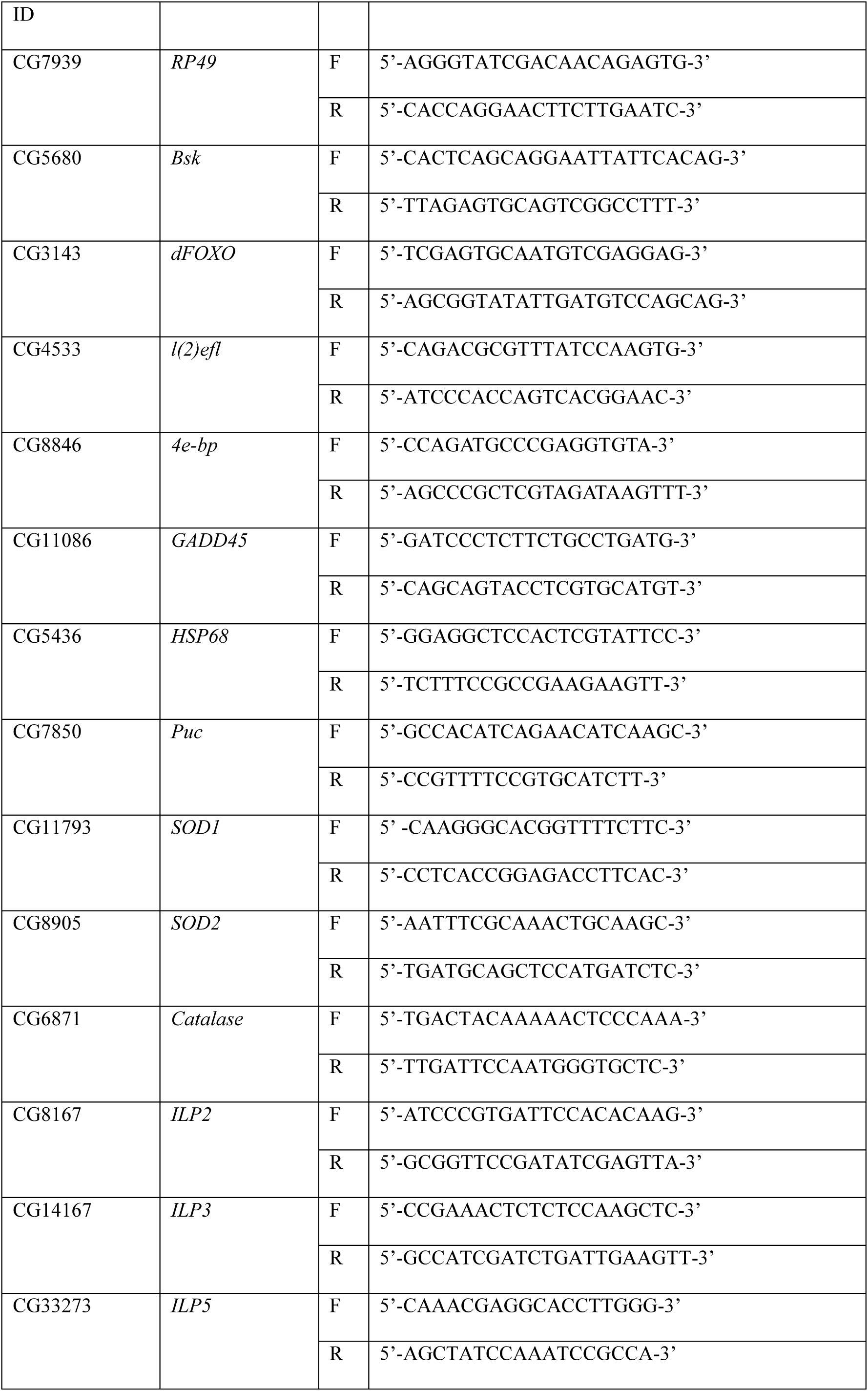
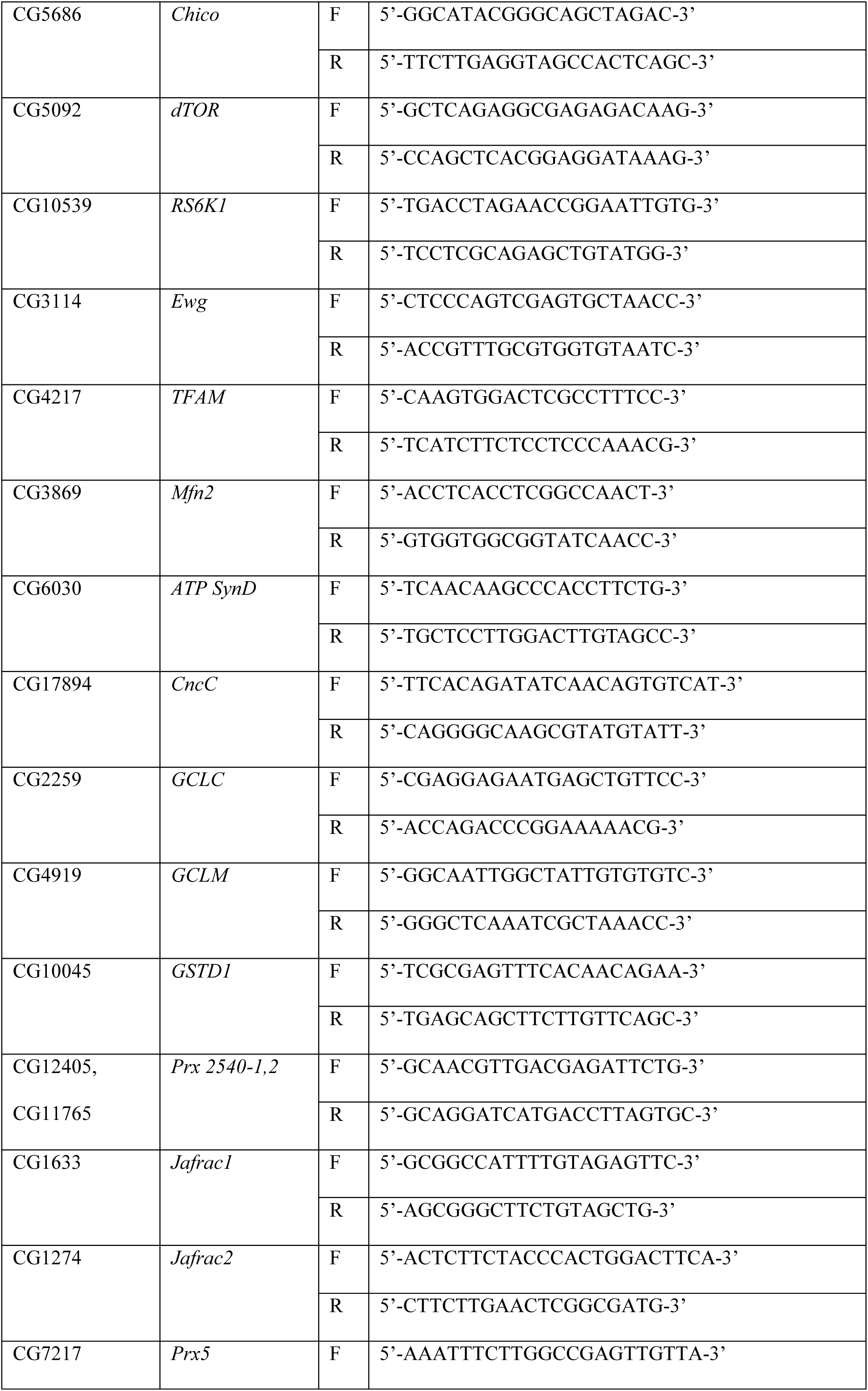
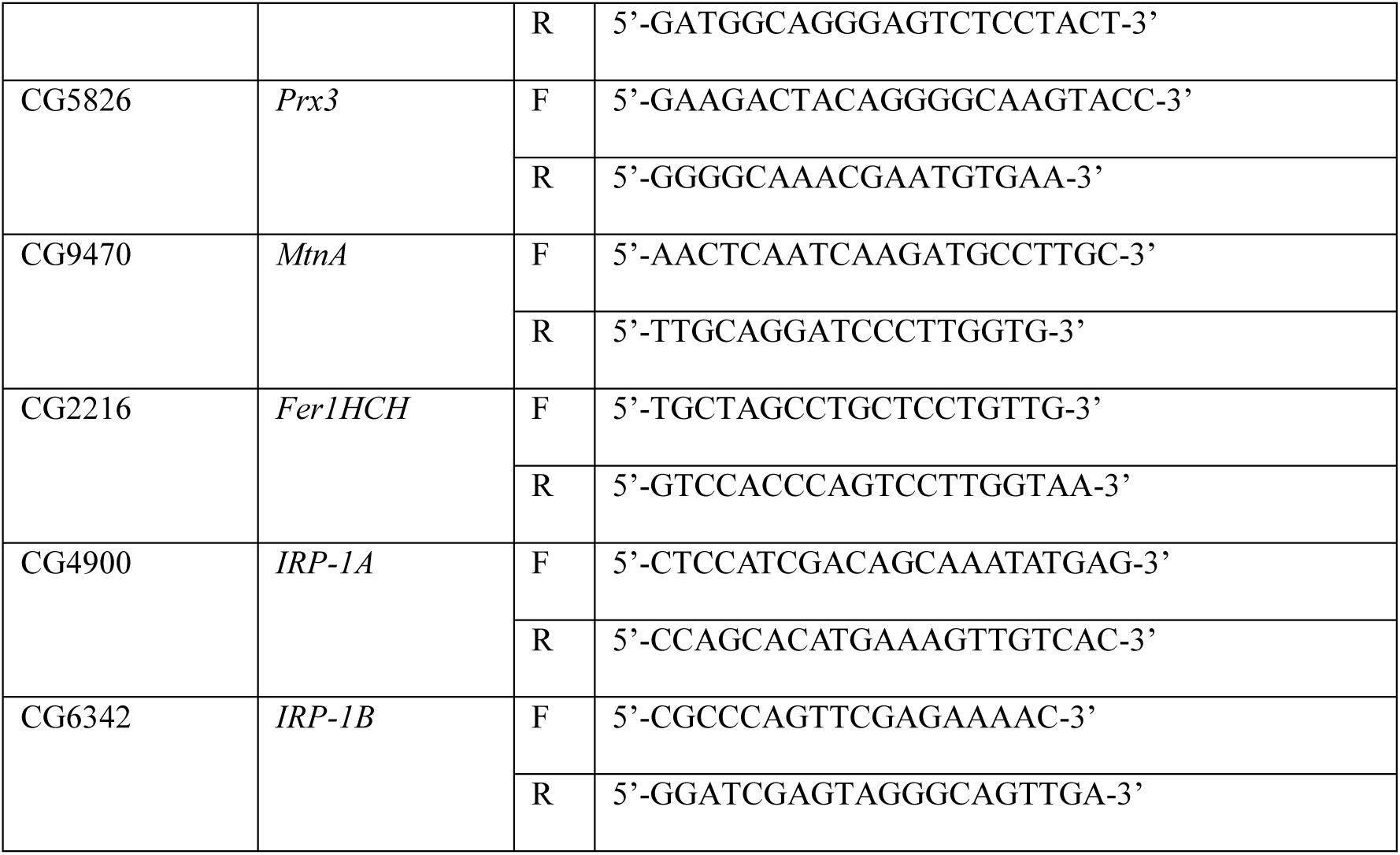
Details of the gene specific primers for expression analysis

The specificity of the primers were validated with melting curve analysis (**Supplementary Information 2).**

#### 2.2.5. Statistical analysis

Graphs were created using GraphPad Prism 8.4.3 software, statistical analysis was completed, and results were expressed as the mean ± standard error of the mean (SEM). A one-way analysis of variance (ANOVA) followed by Tukey post-hoc test and unpaired t-test was carried out to draw significance for the gene expression data. P-values under 0.05 were regarded as significant.

## 3. Results

### 3.1. CU-mediated ALSS modulation of *JNK/Bsk* Signalling pathway

#### 3.1.1. CU fails to rescue diminished *Bsk* expression in the PD brain during HP and TP

*Bsk-*signalling promote neuroprotection when activated in moderation (Gan et al., 2021). In the present study, *Bsk* expression in the PD brain was inhibited by 39% (P<0.01) during HP and by 51% (P<0.001) during TP. CU intervention did not alter the diminished expression of *Bsk* during both the adult life stages. However, feeding of CU *per se* upregulated *Bsk* expression by 87% (Compared to the control brain) (P<0.001) during HP but not during TP (**Fig 1a**). The observation suggests that while CU can modulate *Bsk* expression during HP in the physiologic condition, its intervention has no influence in modulating *Bsk* in PD brain of both life phases. Further, we tried to investigate the aging-associated changes in *Bsk* expression in the healthy brain. It was observed that in the TP fly brain, *Bsk* expression is downregulated by 54% (P<0.001) as compared to that of the HP (**Fig S1a**).

**Figure 1:**
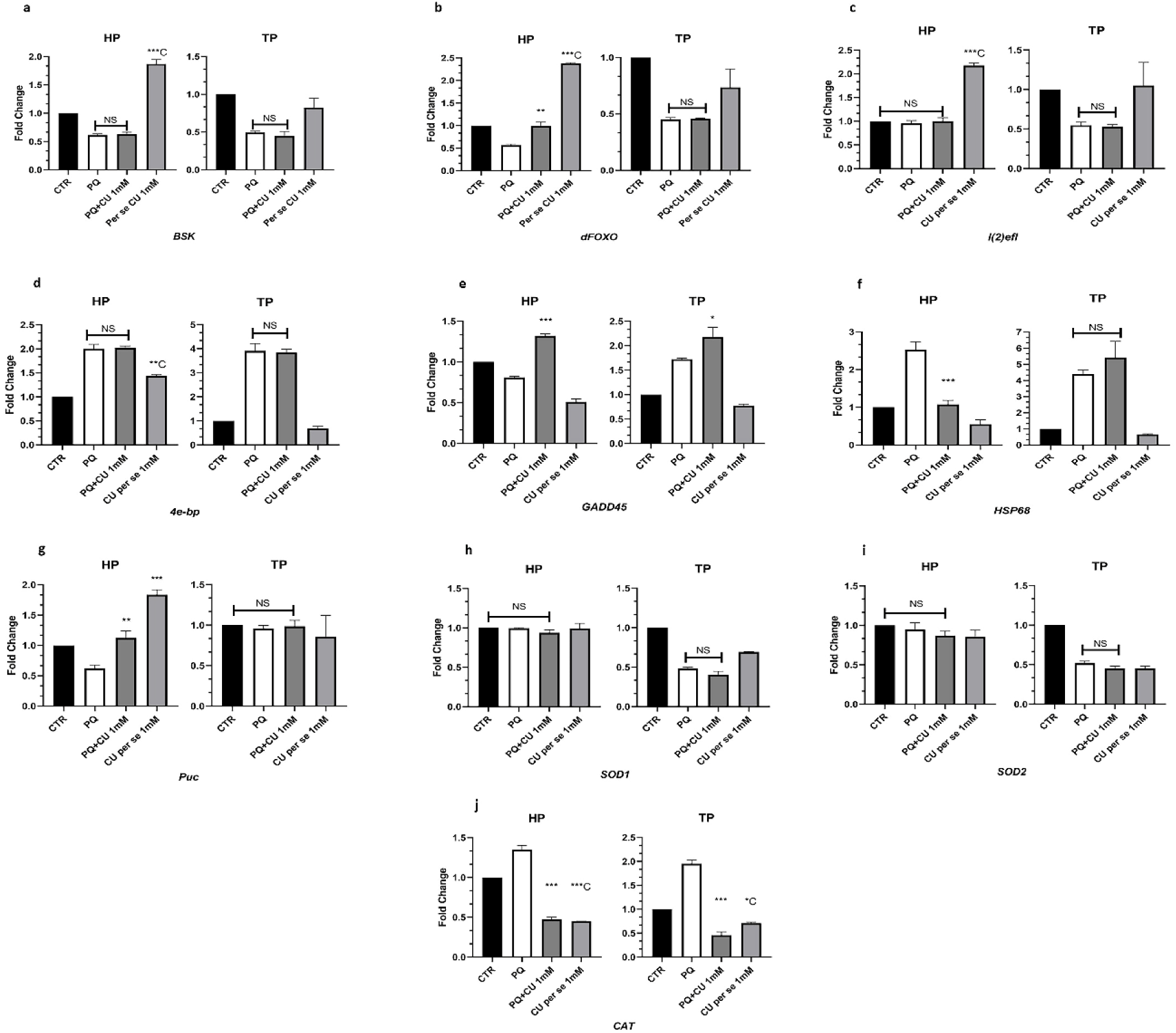
CU-mediated ALSS differential expression of molecular players involved in *JNK/Bsk* mediated stress response pathway in *Drosophila* model of PD. In the PD brain *Bsk* and *dFOXO* expression was inhibited during both HP and TP. CU intervention differentially regulates *Bsk-dFOXO* axis-mediated stress response through the rescue of diminished *dFOXO* only during HP (a,b). However, with CU intervention expression of *dFOXO* downstream targets *l(2)efl* (Unaltered during HP and inhibited during TP upon PQ treatment) and *4e-bp* (Upregulated upon PQ treatment in both HP and TP) remain unaltered (Compared to PD brain) during both the adult life stages (c,d). CU intervention mediated HP-specific rescue of upstream *dFOXO*, resulted in rescue of diminished *GADD45* level (Compared to PD brain) during HP, whereas during TP expression of the same is further upregulated (Compared to PD brain) with CU intervention (e). Further, CU-intervention differentially modulates *Bsk* downstream *HSP68* by inhibiting the upregulation (Compared to PD brain) during HP but not during TP (f). Similarly, CU intervention also differentially modulates *Bsk* downstream *Puc* (Inhibited during HP and unaltered during TP upon PQ treatment) through the rescue of diminished *Puc* level (Compared to PD brain) during HP (g). On the other hand, expression of *dFOXO* downstream antioxidant genes *SOD1* and *SOD2* (Unaltered during HP and inhibited during TP upon PQ treatment) were unaltered (Compared to PD brain) with CU intervention during both the adult life stages, whereas expression of *CAT* (Upregulated upon PQ treatment during both HP and TP) was inhibited with CU intervention during both the adult life stages (h,i,j). Overall insights suggest that CU-mediated ALSS neuroprotection underlies differential modulation of *dFOXO*, *dFOXO* driven *GADD45*, *HSP68* and *Puc* of the *Bsk* signalling pathway. Significance was drawn by analysing the data of a minimum of three replicates with one-way ANOVA followed by Tukey post hoc test. [*p<0.05; **p<0.01; ***p<0.001; NS: Not significant - compared to PQ treated group], [*C p<0.05; **C p<0.01; ***C p<0.001 - compared to Control (CTR) group].

#### 3.1.2. *dFOXO* level was diminished in the PD brain and CU intervention rescues only during HP

The downstream effect of *Bsk* signalling is brought about activation of transcription factor *dFOXO*, promoting gene expressions associated to adaptive stress response, neuroprotection, neuronal maintenance and longevity [25,43,44]. In the present study, *dFOXO* expression in the PD brain was inhibited by 44% (P<0.01) during HP and by 55% (P<0.01) during TP (**Fig 1b**). However, CU intervention (500 µM and 1 mM) rescued diminished *dFOXO* level significantly (P<0.05 and P<0.01) during HP but not during TP. CU *per se* feeding also upregulated *dFOXO* expression by 2 fold (compared to control brain) (P<0.001) during HP but not during TP (**Fig 1b**). The observation suggests that *dFOXO* expression modulation in PD and physiologic conditions is a feature of CU that is restricted only to HP. Hence, CU-mediated ALSS neuroprotection underlies the modulation of *dFOXO* of the *Bsk* signalling pathway. Further investigation revealed that in healthy TP brain, *dFOXO* expression is downregulated by 16% (P<0.01) as compared to that of HP (**Fig S1b**).

#### 3.1.3. *l(2)efl* expression in the PD brain was diminished only during TP and CU fails to alter the same

*l(2)efl* is regulated by *dFOXO* signalling and codes for an sHSP (small heat shock protein) of the α-crystallin family in *Drosophila* and is actively involved in promoting neuroprotection and longevity [28,45]. In the present study, *l(2)efl* expression in the PD brain was unaltered during HP but inhibited by 46% (P<0.05) during TP (**Fig 1c**). CU intervention did not alter the diminished expression of *l(2)efl* during TP. However, CU *per se* feeding upregulated *l(2)efl* expression level by 2 fold (Compared to control brain) (P<0.001) during HP but not during TP (**Fig 1c**). The observation suggests that CU can modulate *l(2)efl* expression only during the physiologic condition of HP. Further *l(2)efl* expression did not alter with age (**Fig S1c**).

#### 3.1.4. CU has no influence on upregulated *4e-bp* expression in the PD brain

*dFOXO* signalling regulated *4e-bp* in *Drosophila* codes for a protein that inhibits translation under stress conditions, thereby indirectly preventing protein aggregation and energy consumption [28, 46]. In the present study, *4e-bp* expression in the PD brain was upregulated by 2 fold (P<0.001) during HP and by 4 fold (P<0.001) during TP (**Fig 1d**). CU intervention did not alter the upregulated expression of *4e-bp* during both adult life stages. However, CU *per se* feeding upregulated *4e-bp* expression level by 44% (Compared to the control brain) (P<0.01) during HP but not during TP (**Fig 1d**). The observation suggests that CU can modulate *4e-bp* expression only during the physiologic condition of HP. Further, *4e-bp* expression was also upregulated with aging [58% (P<0.001) higher in TP compared to HP] (**Fig S1d**).

#### 3.1.5. CU rescues diminished *GADD45* level in the PD brain only during HP

*dFOXO/FOXO* signalling regulated *GADD45* modulates DNA damage repair and can promote neuroprotection and apoptosis in *Drosophila* and *in-vitro* neuronal cells respectively [47–51]. In the present study, *GADD45* expression in the PD brain was inhibited by 20% (P<0.01) during HP, but upregulated by 72% (P<0.01) during TP (**Fig 1e**). However, CU intervention rescued the diminished *GADD45* level significantly (P<0.001) during HP, and it further enhanced (P<0.05) the upregulated *GADD45* during TP (**Fig 1e**). The observation suggests that CU-mediated ALSS neuroprotection underlies the rescue of diminished *GADD45* level during HP. Further investigation revealed that *GADD45* expression is upregulated with aging [3 fold (P<0.001) higher in TP compared to HP] (**Fig S1e**).

#### 3.1.6. *HSP68* was upregulated in the PD brain but CU rescues only during HP

*Bsk/JNK* signalling regulates *HSP* and *HSP68* may not be different. Its molecular chaperone-mediated response promotes longevity and stress resistance [27,29,52]. In the present study, *HSP68* expression in the PD brain was upregulated by 2.5 fold (P<0.001) during HP and by 4 fold (P<0.01) during TP (**Fig 1f**). However, CU intervention significantly inhibited (P<0.001) the upregulated *HSP68* level during HP but not during TP. The observation suggests that CU intervention can inhibit proteinopathic stress and molecular dysregulations associated with NDD. Therefore, CU-mediated ALSS neuroprotection also involves the rescue of altered *HSP68* level. The investigation also revealed *HSP68* expression is upregulated with aging [2 fold (P<0.05) higher in TP compared to HP] (**Fig S1f**).

#### 3.1.7. *Puc* expression diminished in the PD brain only during HP and CU rescues

Hyper-activation of *Bsk* signalling is prevented by the expression of Bsk-phosphatase *Puc* through a negative feedback loop [29,53–55]. In the present study, *Puc* expression in the PD brain was inhibited by 38% (P<0.05) during HP, but expression of the same remained unaltered during TP. However, CU intervention rescued the diminished *Puc* level significantly (P<0.01) during HP (**Fig 1g**). Further, CU *per se* feeding also upregulated *Puc* expression by 83% (Compared to the control brain) (P<0.001) during HP but not during TP (**Fig 1g**). The observation suggests that *Puc* expression modulation in PD and physiologic conditions is a feature of CU, that is restricted only to HP. Therefore, CU-mediated ALSS neuroprotection underlies the modulation of *Puc*. Investigation revealed that *Puc* expression is downregulated with aging [51% (P<0.001) lower in TP compared to HP] (**Fig S1g**).

#### 3.1.8. *SOD1* and *SOD2* diminished only during TP and CU fails to rescue

*dFOXO/FOXO* signalling regulated *SOD1* and *SOD2* neutralizes superoxide in cytosol and mitochondria respectively and promotes neuroprotection and longevity [56–61]. In the present study, *SOD1* and *SOD2* expression in the PD brain was unaltered during HP but were inhibited by 52% (P<0.001) and 49% (P<0.001) respectively during TP (**Fig 1 h,i**). CU intervention did not alter the diminished *SOD1* and *SOD2* levels during TP. The observation suggests that CU intervention has no role in modulating *SOD1* and *SOD2* in PD. It was observed that, while *SOD1* expression is unaltered with aging and *SOD2* expression is downregulated in the TP brain by 40% (P<0.01) compared to HP (**Fig S1 h,i**).

#### 3.1.9. CU inhibits upregulated *CAT* in the PD brain during both HP and TP

*dFOXO/FOXO* signalling regulated *CAT* expression is induced under stress as defense mechanism against peroxide and it also promotes longevity [56,57,59–63]. In the current study, *CAT* expression in the PD brain was upregulated by 35% (P<0.001) during HP and by nearly 2 fold (P<0.001) during TP (**Fig 1j**). However, CU intervention significantly inhibited (P<0.001) the upregulated *CAT* level during both the adult life stages. Further, feeding of CU *per se* inhibited *CAT* expression by 56% (P<0.001) during HP and by 30% (P<0.05) during TP (Compared to the control brain) (**Fig 1j**). The observation suggests that *CAT* expression modulation in PD and physiologic conditions is a feature of CU and is not restricted to the adult life stages. The ability of CU intervention to inhibit *CAT* upregulation in the PD brain signifies inhibition of peroxide stress, but TP flies are not rescued with CU intervention. Therefore, modulation of *CAT* may be necessary, but is not the critical to ALSS neuroprotection. It was observed that *CAT* expression in aging brain is unaltered (**Fig S1j**).

### 3.2. CU-mediated ALSS modulation of *IIS-dTOR* Signalling pathway

#### 3.2.1. *ILP2*, *ILP5* expression diminished in the PD brain while *ILP3* was unaltered, but CU inhibits all the three *ILP* during HP and TP

*Drosophila ILP*s are analogous to mammalian insulin/insulin-like signalling pathway (*IIS*), mediating anabolic processes and ablation of *IIS* components extends lifespan and enhance stress resistance [31,64–67]. In the present study, *ILP2* expression in the PD brain was inhibited by 19% (P<0.001) during HP and by 17% (P<0.01) during TP. Diminished expression in the PD brain was also observed for the *ILP5* which was inhibited by 31% (P<0.001) during HP and by 29% (P<0.01) during TP. *ILP3* expression however was not altered in the PD brain of both life phases (**Fig 2 a,b,c**). CU intervention further diminished the expression of *ILP2* and *ILP5* during HP (P<0.05 and P<0.01 respectively) and TP (P<0.01 and P<0.05 respectively). CU intervention also inhibited *ILP3* expression during HP (P<0.01) and TP (P<0.001). The observation suggests that although only *ILP2* and *ILP5* are suppressed in the PD brain, CU intervention may promote suppression of *IIS* through the inhibition of all three *ILP*s. Inhibition of *ILPs* is therapeutic to organisms during stress, but transition phase flies are not rescued with CU intervention. Therefore, *ILP*s modulation in the PD brain may be a common mode of action of CU intervention and is not critically involved in ALSS neuroprotection. It is revealed that *ILP2* expression was enhanced by 30% (P<0.001), *ILP3* expression was diminished by 43% (P<0.01) and *ILP5* expression was not altered with aging (Compared to HP) (**Fig S2 a,b,c**).

**Figure 2:**
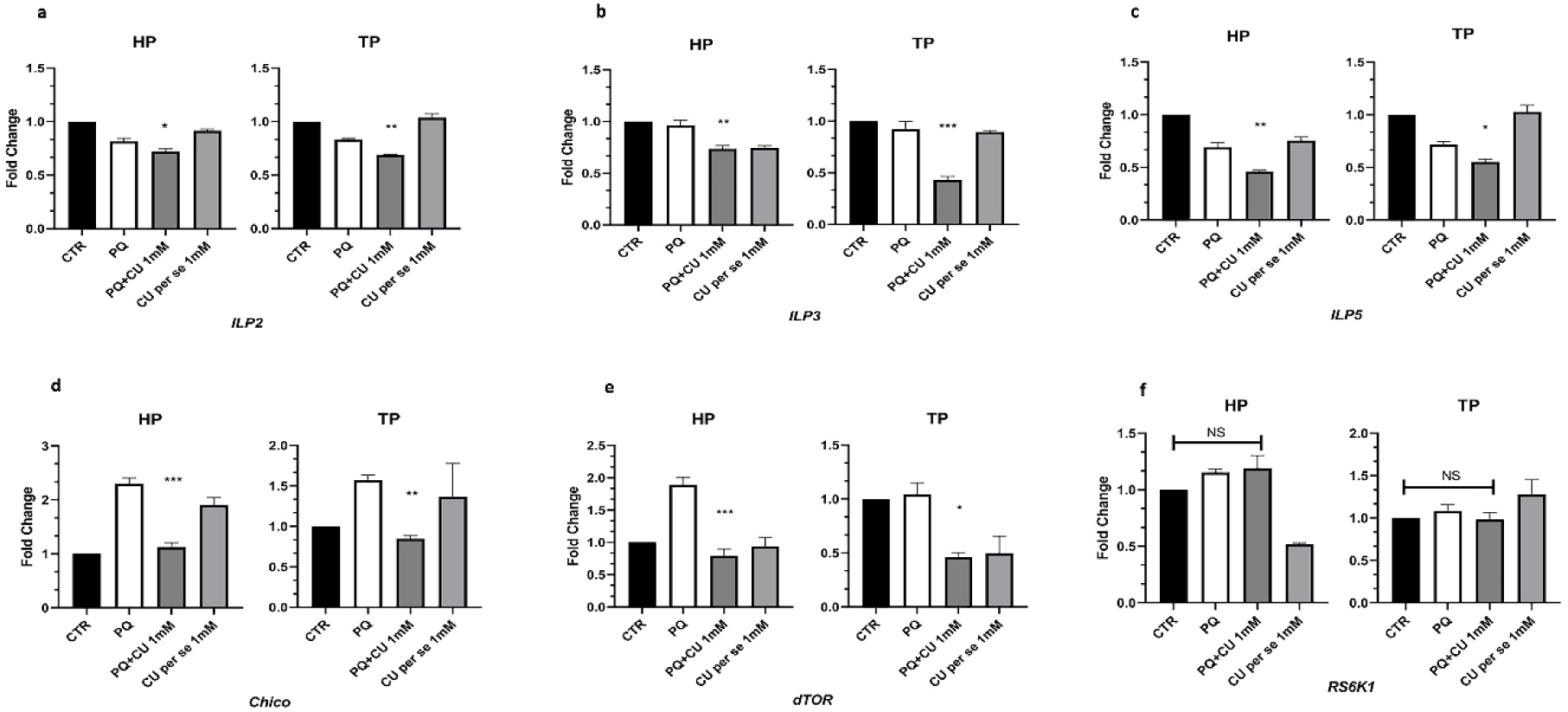
CU-mediated ALSS differential expression of molecular players involved in *IIS-dTOR* mediated anabolic pathway in *Drosophila* model of PD. In the PD brain *ILP2* and *ILP5* expression was inhibited, whereas *ILP3* expression remain unaltered during HP and TP. CU intervention inhibits *IIS* signaling by further inhibiting *ILP2*, *ILP5* and inhibiting *ILP3* during both adult life stages (a,b,c). Similarly, CU intervention inhibits *IIS* signaling by inhibiting *Chico* (Upregulated during HP and TP upon PQ treatment) (d). Also, CU intervention inhibits *IIS* downstream *dTOR* (Upregulated during HP and unaltered during TP upon PQ treatment) during both the adult life stages (e). But PD condition and CU intervention did not alter *dTOR* downstream *RS6K1* expression (f). Overall insights suggest that CU intervention inhibits *IIS- dTOR* signaling during both the adult life stages. Though inhibition of *IIS-dTOR* inhibition is protective, TP flies are not rescued with CU intervention. Therefore, modulation of *IIS-dTOR* signaling is not critically involved in CU-mediated ALSS neuroprotection. Significance was drawn by analysing the data of a minimum of three replicates with one-way ANOVA followed by Tukey post hoc test. [*p<0.05; **p<0.01; ***p<0.001; NS: Not significant - compared to PQ treated group].

#### 3.2.2. CU rescues upregulated *Chico* in the PD brain during HP and TP

*Drosophila Chico* codes for mammalian analogous the insulin receptor substrate for the propagation of *IIS* and inhibition of *Chico* also promotes life span extension cum stress resistance [31,64,68,69]. In the present study, *Chico* expression in the PD brain was upregulated by 2 fold (P<0.001) during HP and by 56% (P<0.05) during TP (**Fig 2d**). However, CU intervention significantly inhibited the upregulation during both the HP (P<0.001) and TP (P<0.01) (**Fig 2d**). The observation suggests that in both the adult life stages enhanced *IIS* signalling in PD condition is inhibited by CU intervention through inhibition of upregulated *Chico*. Inhibition of *Chico* is therapeutic to organisms under stress conditions, but TP flies are not rescued with CU intervention. Therefore, *Chico* modulation by CU intervention is not critically associated with ALSS neuroprotection. It was observed that natural aging enhances brain *Chico* level [2 fold (P<0.01) higher in TP compared to HP] (**Fig S2d**).

#### 3.2.3. PD upregulated *dTOR* during HP and CU rescues, whereas in the TP PD brain *dTOR* expression was unaltered but CU inhibits the same

*IIS* signalling-mediated activation of *dTOR* signalling ensures anabolic functions and reduced *dTOR* signalling promotes life span extension cum stress resistance [31,32,64,70]. In the present study, *dTOR* expression in the PD brain was enhanced by 88% (P<0.001) during HP, whereas no alteration was observed during TP (**Fig 2e**). CU intervention significantly inhibited the upregulation during HP (P<0.001) and inhibited *dTOR* level (P<0.05) during TP (**Fig 2e**). The observation suggests that CU intervention inhibits *dTOR* signalling in PD brain. Inhibition of *dTOR* is therapeutic under stress conditions, but TP flies are not rescued with CU intervention. Therefore, *dTOR* modulation in the PD brain by CU is not critically associated with ALSS neuroprotection. It was observed that *dTOR* level in the brain is diminished with aging [21% (P<0.05) lesser in TP compared to HP] (**Fig S2e**).

#### 3.2.4. *RS6K1* expression was not altered in the PD brain and with CU intervention

*dTOR* signalling also involves the activation of kinase coded by *RS6K1* and similarly, its inhibition promotes life span extension cum stress resistance [31,32,64,70]. In the present study, *RS6K1* expression was not altered in the PD condition or with CU intervention (**Fig 2f**). The observation suggests that CU-mediated ALSS neuroprotection does not involve *RS6K1* modulation. It was observed that *RS6K1* expression is diminished with aging [64% (P<0.001) lesser in TP compared to HP (**Fig S2f**)].

### 3.3. CU-mediated ALSS modulation of components involved in mitochondrial dynamics

#### 3.3.1. CU fails to rescue diminished *Ewg* expression in the PD brain during HP and TP

*IIS-dTOR/mTOR*-mediated anabolic processes initiate mito-biogenesis through transcription factor *NRF1/Ewg* [71,72]. In the present study, *Ewg* expression in the PD brain was inhibited by 38% (P<0.01) during HP and by 24% (P<0.05) during TP (**Fig 3a**). CU intervention did not alter the diminished expression of *Ewg* during both the adult life stages (**Fig 3a**). The observation suggests that CU intervention has no influence in modulating *Ewg* in the PD brain of both adult life stages. Therefore, CU-mediated ALSS neuroprotection do not underlie *Ewg* modulation. The investigation also revealed that *Ewg* expression is inhibited with natural aging [25% (P<0.001) lesser in TP compared to HP] (**Fig S3a**).

**Figure 3:**
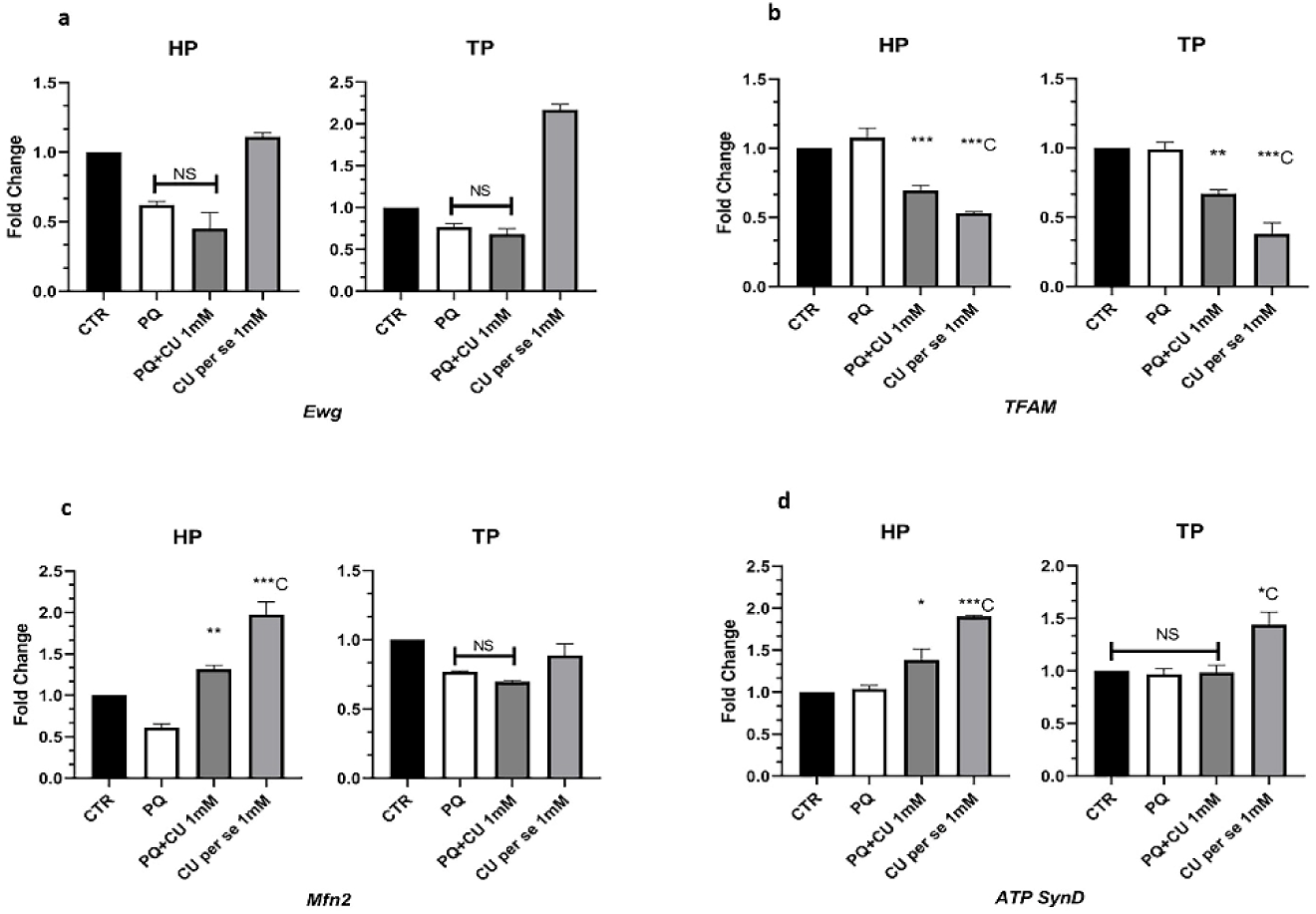
CU-mediated ALSS differential expression of molecular players involved in mitochondrial dynamics in the *Drosophila* model of PD. In the PD brain mito-biogenesis mediating transcription factor *Ewg* expression was inhibited during both HP and TP and CU intervention fails to alter the diminished *Ewg* level (a). However, with CU intervention *Ewg* downstream *TFAM* level (Unaltered upon PQ treatment) was inhibited during both HP and TP (b). In the PD brain *Mfn2* level was inhibited during both HP and TP. CU differentially regulates mitochondrial quality control through the rescue of diminished *Mfn2* level during HP, but not during TP (c). Similarly, CU intervention differentially regulates mitochondrial respiratory capacity, through upregulating *ATP SynD* (Unaltered upon PQ treatment) level only during HP (d). Significance was drawn by analysing the data of a minimum of three replicates with one-way ANOVA followed by Tukey post hoc test. [*p<0.05; **p<0.01; ***p<0.001; NS: Not significant - compared to PQ treated group], [*C p<0.05; **C p<0.01; ***C p<0.001 - compared to Control (CTR) group].

#### 3.3.2. *TFAM* expression was unaltered in the PD brain, but CU intervention inhibits during HP and TP

*Ewg* signalling regulates *TFAM* mediating mtDNA replication for mito-biogenesis (71- 73). In the present study, *TFAM* expression in the PD brain was not altered during HP and TP (**Fig 3b**). However, CU intervention significantly inhibited *TFAM* level during HP (P<0.001) and TP (P<0.01). Further, feeding of CU *per se* inhibited *TFAM* expression by 47% (P<0.001) during HP and by 62% (P<0.001) during TP (**Fig 3b**). The observation suggests that *TFAM* modulation in PD and physiologic conditions is a feature of CU, which is common to both the adult life stages. *TFAM* inhibition may be protective, but TP flies are not rescued with CU intervention. Therefore, modulation of *TFAM* is not critically involved in CU-mediated ALSS neuroprotection. The investigation also revealed that *TFAM* expression is diminished with natural aging [30% (P<0.001) lesser in TP compared to HP] (**Fig S3b**).

#### 3.3.3. *Mfn2* level was diminished in the PD brain and CU intervention rescues only during HP

*dFOXO/*FOXO signalling regulates *Mfn2* to mediate mitochondrial quality control, complementation and it actively promotes neuroprotection [33,34,74–76]. In the present study, *Mfn2* expression in the PD brain was inhibited by 39% (P<0.05) during HP and by 24% (P<0.05) during TP (**Fig 3c**). However, CU intervention rescued diminished *Mfn2* level significantly (P<0.01) only during HP but not during TP. Feeding of CU *per se* also upregulated *Mfn2* expression by nearly 2 folds (P<0.001) only during HP (**Fig 3c**). The observation suggests that *Mfn2* expression modulation in PD and physiologic conditions is a feature of CU that is restricted only to HP. Therefore, CU mediated ALSS neuroprotection underlies modulation of *Mfn2*. It was found that *Mfn2* expression is enhanced with aging [20% (P<0.01) higher in TP compared to HP] (**Fig S3c**).

#### 3.3.4. *ATP SynD* expression was unaltered in the PD brain but CU intervention upregulates only in HP

*ATP SynD* codes for the ATP synthase subunit associated with the Complex V (Cl-V) of the mitochondrial respiratory chain [77]. In the present study, *ATP SynD* expression in the PD brain was not altered during HP and TP. CU intervention upregulated *ATP SynD* expression (P<0.05) only during HP only but not during TP (**Fig 3d**). Also, feeding of CU *per se* upregulated *ATP SynD* expression by nearly 2 folds (P<0.001) during HP and by 43% (P<0.001) during TP (**Fig 3d**). The observation suggests that CU-mediated modulation of *ATP SynD* under the PD condition is restricted only to HP. Further, the PD condition might not specifically affect Cl-V of the respiratory chain, but CU-mediated differential modulation of *ATP SynD* may contribute to better respiratory health during HP. It was observed that *ATP SynD* downregulates with natural aging [70% (P<0.001) lesser in TP compared to HP] (**Fig S3d**).

### 3.4. CU-mediated ALSS modulation of phase II ADS

#### 3.4.1. *CncC* was diminished in the PD brain and CU intervention rescues only during HP

*CncC* (Mammalian *Nrf2*) actively participates in the regulation of phase II ADS and is a potential target of neuroprotection in PD [78–80]. In the present study, *CncC* expression in the PD brain was inhibited by 55% (P<0.001) during HP and by 51% (P<0.01) during TP (**Fig 4a**). However, CU intervention rescued (P<0.01) diminished *CncC* level only during HP but not during TP (**Fig 4a**). The observation suggests that CU-mediated ALSS neuroprotection underlies HP-specific rescue of *CncC*. Further, the investigation revealed that *CncC* expression is upregulated with natural aging [2 fold (P<0.001) higher in TP compared to HP] (**Fig S4a**).

**Figure 4:**
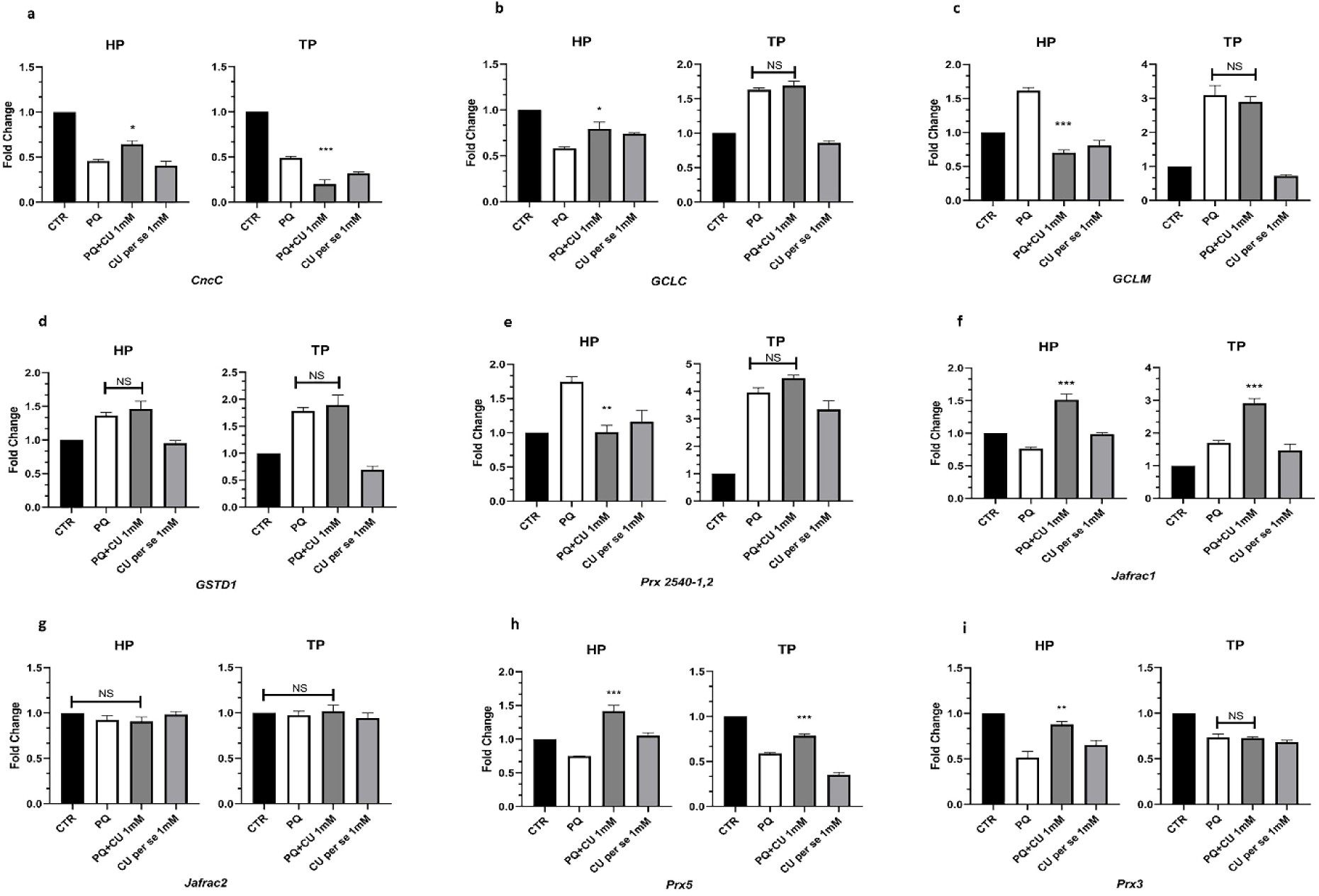
CU-mediated ALSS differential expression of molecular players involved in phase II ADS in the *Drosophila* model of PD. In the PD brain *CncC* expression was inhibited during both HP and TP. CU intervention differentially regulates phase II ADS through the rescue of diminished *CncC* only during HP (a). CU intervention mediated HP-specific rescue of upstream *CncC* resulted in the rescue of diminished *GCLC* level (Compared to PD brain) during HP, whereas during TP expression of upregulated *GCLC* was not altered with CU intervention (b). Further, CU-intervention differentially modulates *CncC* downstream *GCLM* by inhibiting the upregulation (Compared to PD brain) during HP but not during TP (c) CU intervention did not alter CncC downstream *GSTD1* (Upregulated upon PQ treatment during HP and TP) expression level (d). Further, CU-intervention differentially modulates *CncC* downstream *Prx 2540-1,2* by inhibiting the upregulation (Compared to PD brain) during HP but not during TP (e). CU intervention mediated HP-specific rescue of upstream *CncC* resulted in the rescue of diminished *Jafrac1* level (Compared to PD brain) during HP, whereas during TP expression of the same was further upregulated with CU intervention (f). However, *Jafrac2* was not altered in the PD brain or with CU intervention (g). The diminished level of *CncC* downstream *Prx5* was rescued (Compared to PD brain) with CU intervention during HP and TP (h). However, the diminished level of *Prx3* was rescued (Compared to PD brain) with CU intervention only during HP, like upstream *CncC* (i). Overall insights suggest that CU-mediated ALSS neuroprotection underlies differential modulation of phase II ADS mediator *CncC*, which results in synchronous differential modulation of *Prx3*. Rescue of diminished *CncC* during HP also results in synchronous rescue of diminished *GCLC* and *Jafrac1*. CU-mediated ALSS neuroprotection also underlies differential modulation of *GCLM* and *Prx 2540-1,2* which is not synchronous to upstream *CncC*. Significance was drawn by analysing the data of a minimum of three replicates with one-way ANOVA followed by Tukey post hoc test. [*p<0.05; **p<0.01; ***p<0.001; NS: Not significant - compared to PQ treated group], [*C p<0.05; **C p<0.01; ***C p<0.001 - compared to Control (CTR) group].

#### 3.4.2. CU rescues diminished *GCLC* level in PD brain only during HP

*CncC*-mediated regulation of phase II ADS and neuroprotection also involves the production and utilization of reduced GSH which is mediated by *GCLC* [78,81,82]. In the present study, *GCLC* expression in the PD brain was inhibited by 42% (P<0.001) during HP, but upregulated by 63% (P<0.001) during TP (**Fig 4b**). However, CU intervention rescued (P<0.05) the diminished *GCLC* level during HP, but failed to alter *GCLC* upregulation during TP (**Fig 4b**). The observation suggests that CU-mediated ALSS neuroprotection underlies the rescue of diminished *GCLC* level during HP. Further investigation revealed that *GCLC* expression is diminished with aging [50% (P<0.001) lesser in TP compared to HP] (**Fig S4b**).

#### 3.4.3. *GCLM* upregulated in PD brain but CU rescues only during HP

*CncC*-mediated regulation of *GCLM* ensures the stability of the GSH synthesis machinery and its gene response is necessary for the prevention of neurodegeneration [82–84]. In the present study, *GCLM* expression in the PD brain was upregulated by 70% (P<0.001) during HP and by 3 fold (P<0.001) during TP (**Fig 4c**). However, CU intervention significantly inhibited (P<0.001) the upregulated *GCLM* level during HP but not during TP (**Fig 4c**). The observation suggests that CU intervention can inhibit some form of stress that is specifically countered by phase II ADS, normalizing the associated molecular dysregulation. Therefore, CU-mediated ALSS neuroprotection involves modulation of *GCLM*. Further, investigation revealed that *GCLM* expression is diminished with aging [25 % (P<0.01) lesser in TP compared to HP] (**Fig S4c**).

#### 3.4.4. CU does not alter the upregulated *GSTD1* expression in PD brain

*CncC* and *Bsk* signalling regulated *GSTD1* mediates toxin neutralization through reduced and it is often upregulated in the presence of toxins cum xenobiotics [29,62,80,85–88]. In the present study, *GSTD1* expression in PD brain was upregulated by 36% (P<0.01) during HP and by 80% (P<0.01) during TP (**Fig 4d**). CU intervention did not alter the upregulated expression of *GSTD1* during both the adult life stages. The observation suggests that CU do not modulate *GSTD1* expression in the PD brain. Therefore, CU-mediated ALSS neuroprotection does not underlie *GSTD1* modulation. Further, *GSTD1* expression was also enhanced with aging [80% (P<0.001) higher in TP compared to HP] (**Fig S4d**).

#### 3.4.5. *Prx 2540-1,2* was upregulated in the PD brain but CU rescues only during HP

*Prx 2540-1,2* is a copy variant of the *Prx 2540* and may be regulated by *CncC* signalling and it also mediates pro-inflammatory pathways [89–91]. In the present study, *Prx 2540- 1,2* expression in PD was upregulated by 70% (P<0.01) during HP and by 4 fold (P<0.001) during TP (**Fig 4e**). However, CU intervention significantly inhibited (P<0.01) the upregulated *Prx 2540-1,2* level only during HP but not during TP (**Fig 4e**). The observation suggests that CU intervention can inhibit some form of stress that is specifically countered by phase II ADS and/or inhibit pro-inflammatory signature by inhibiting associated molecular dysregulation. Therefore, CU-mediated ALSS neuroprotection involves modulation of *Prx 2540-1,2* expression. Further, investigation also revealed that *Prx 2540-1,2* expression is upregulated with aging [2 fold (P<0.01) higher in TP compared to HP] (**Fig S4e**).

#### 3.4.6. CU rescues diminished *Jafrac1* level in the PD brain only during HP

*Jafrac1* is regulated under *CncC* and *Bsk-dFOXO* signalling and is actively involved in neuroprotection and stress resistance [90,92,93]. In the present study, *Jafrac1* expression in PD brain was inhibited by 24% (P<0.05) during HP, but upregulated by 70% (P<0.05) during TP (**Fig 4f**). However, CU intervention rescued diminished *Jafrac1* level significantly (P<0.001) during HP, but during TP the upregulation of *Jafrac1* was further enhanced (P<0.001) (**Fig 4f**). The observation suggests that CU-mediated ALSS neuroprotection underlies the rescue of diminished *Jafrac1* level during HP. Further, the investigation revealed that *Jafrac1* expression is diminished with aging [56% (P<0.001) lesser in TP compared to HP] (**Fig S4f**).

#### 3.4.7. *Jafrac2* expression was not altered in the PD brain and with CU intervention

*Jafrac2* is regulated under *CncC* signalling and promotes longevity, stress resistance and also promotes apoptosis depending on its signalling intensity [90,94]. In the present study, *Jafrac2* expression was not altered in PD brain and with CU intervention, during both the adult life stages (**Fig 4g**). No alteration in expression was also observed upon CU *per se* feeding during both the adult life stages (**Fig 4g**). The observation suggests that CU-mediated ALSS neuroprotection does not underlie *Jafrac2* modulation. *Jafrac2* expression is upregulated with aging [68% (P<0.001) higher in TP compared to HP] (**Fig S4g**).

#### 3.4.8. CU rescues diminished *Prx5* expression in the PD brain during HP and TP

*CncC*-regulated *Prx5* codes for mitochondria and cytosol-specific peroxiredoxin that extensively promotes longevity, apoptosis prevention and anti-inflammatory response [90, 95–97]. In the present study, *Prx5* expression in the PD brain was inhibited by 26% (P<0.05) during HP and by 42% (P<0.001) during TP (**Fig 4h**). However, CU intervention rescued diminished *Prx5* level significantly (P<0.001) during both the adult life stages (**Fig 4h**). But TP flies are not rescued with CU intervention. Therefore, *Prx5* modulation by CU in the PD brain may be important but is not critical to ALSS neuroprotection. Further investigation revealed that *Prx5* expression is upregulated with aging [25% (P<0.01) higher in TP compared to HP] (**Fig S4h**).

#### 3.4.9. *Prx3* level was diminished in the PD brain and CU intervention rescues only during HP

*CncC* and *dFOXO/FOXO* regulated *Prx3* codes for only mitochondria-specific peroxiredoxin that promote longevity, stress resistance cum neuroprotection [49,90,96]. In the present study, *Prx3* expression in the PD brain was inhibited by 50% (P<0.001) during HP and by 30% (P<0.001) during TP (**Fig 4i**). However, CU intervention rescued (P<0.01) diminished *Prx3* level only during HP but not during TP (**Fig 4i**). The observation suggests that CU-mediated ALSS neuroprotection underlies the modulation of *Prx3.* Further investigation also revealed that *Prx3* expression is upregulated with aging [30% (P<0.001) higher in TP compared to HP] (**Fig S4i**).

### 3.5. CU-mediated ALSS modulation of components involved in metal homeostasis

#### 3.5.1. CU partially rescues the upregulated *MtnA* in the PD brain

*MtnA* is responsible for zinc/copper homeostasis that promotes anti-inflammatory, anti- oxidant responses against stress [98,99–102]. In the present study, *MtnA* expression in the PD brain was upregulated by 2 fold (P<0.001) during HP and by nearly 3 fold (P<0.001) during TP (**Fig 5a**). However, CU intervention partially inhibited the *MtnA* upregulation (P<0.001) during both the adult life stages. Also, CU *per se* feeding inhibited *MtnA* expression by 60% (P<0.001) during HP and by 40% (P<0.001) during TP (**Fig 5a**). The observation suggests that *MtnA* expression modulation in the PD and physiologic conditions is a feature of CU, which is common to both the adult the life stages. The ability of CU to partially inhibit *MtnA* upregulation in the PD brain suggests its ability to suppress oxidative stress, inflammation and associated molecular dysregulation. But TP flies are not rescued with CU intervention, suggesting modulation of *MtnA* does not underlie CU-mediated ALSS neuroprotection. Further, *MtnA* expression did not alter with aging (**Fig S5a**).

**Figure 5:**
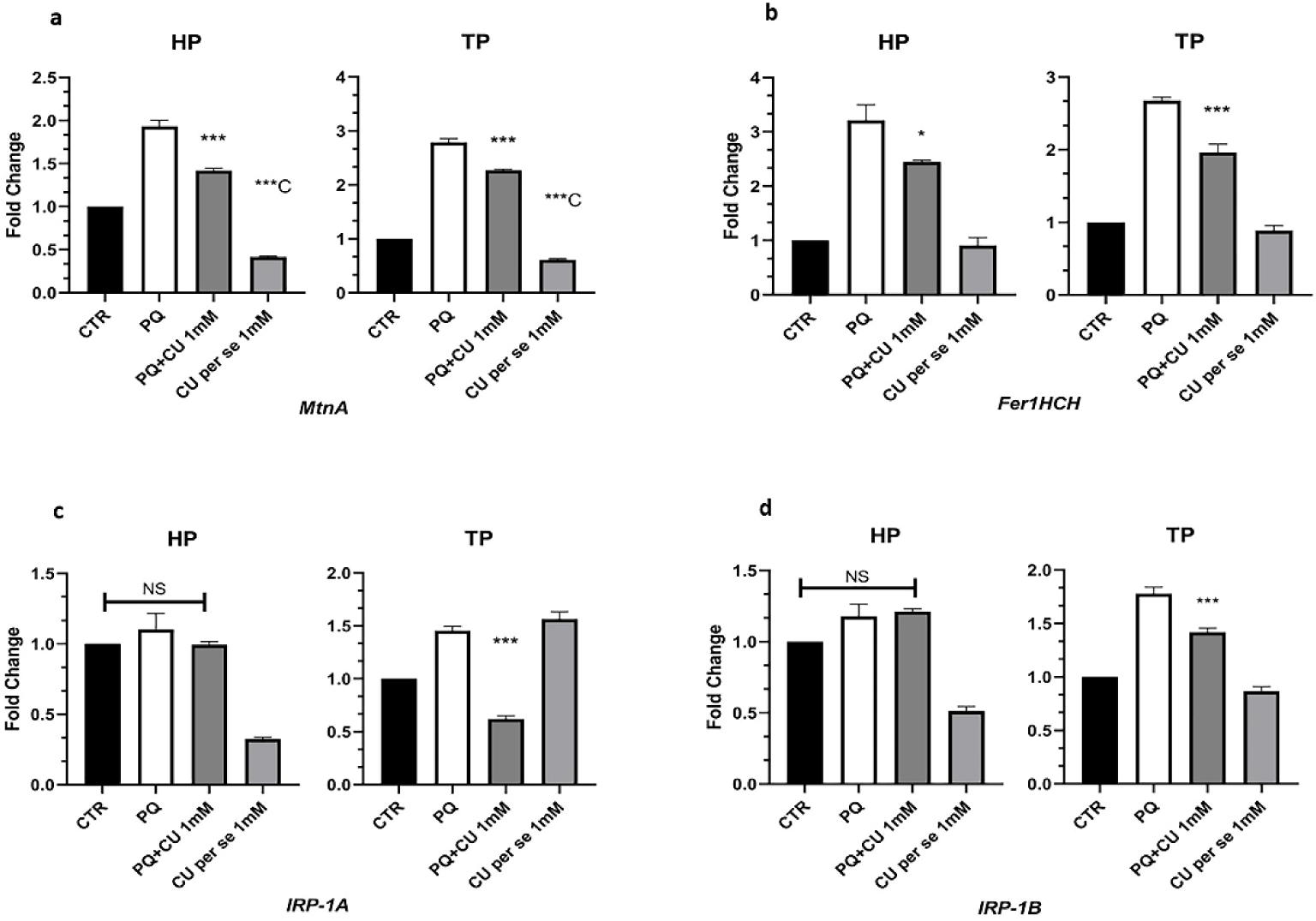
CU-mediated ALSS differential expression of molecular players involved in metal homeostasis in the *Drosophila* model of PD. In the PD brain *MtnA* and *Fer1HCH* expression was upregulated during both the HP and TP. CU intervention partially inhibits the PD-associated upregulation of *MtnA* and *Fer1HCH* during both the adult life stages (a,b). CU intervention in PD brain did not alter *IRP-1A* and *IRP- 1B* (Unaltered during HP and upregulated during TP upon PQ treatment) during HP but during TP it inhibits *IRP* dysregulation by significantly inhibiting *IRP-1A* (Compared to PD brain) and partially inhibiting *IRP-1B* (Compared to PD brain) (c,d). Overall insights suggest that owing to anti-inflammatory, antioxidant and metal-chelating properties CU in the PD brain can partially suppress possible metal (Zinc, copper and iron) imbalance and partially inhibits the dysregulation of *MtnA* and *Fer1HCH*. However, CU cannot fully suppress iron intake during TP as it can only inhibit *IRP-1A* upregulation completely, whereas upregulated *IRP-1B* is inhibited partially in PD brain The. limitation of CU to prevent iron intake in the PD brain during TP highlights its limitation to promote neuroprotection during later adult life stages. Significance was drawn by analysing the data of a minimum of three replicates with one-way ANOVA followed by Tukey post hoc test. [*p<0.05; **p<0.01; ***p<0.001; NS: Not significant - compared to PQ treated group], [*C p<0.05; **C p<0.01; ***C p<0.001 - compared to Control (CTR) group].

#### 3.5.2. CU partially rescues the upregulated *Fer1HCH* in the PD brain

*Fer1HCH* codes for an iron-chelating protein and is necessary for neuronal maintenance cum CNS development in *Drosophila* [103–106]. In the present study, *Fer1HCH* expression in PD brain was upregulated by 3 fold (P<0.001) during HP and by 2.6 fold (P<0.001) during TP (**Fig 5b**). However, CU intervention partially inhibited *Fer1HCH* upregulation during HP (P<0.05) and TP (P<0.001) (**Fig 5b**). The observation suggests the ability of CU to suppress iron dysregulation, and oxidative stress and inhibit associated molecular signature. But TP flies are not rescued with CU intervention, suggesting modulation of *Fer1HCH* does not underlie ALSS neuroprotection. Further, *Fer1HCH* expression was enhanced with aging [2.6 folds (P<0.001) higher in TP compared to HP] (**Fig S5b**).

#### 3.5.3. *IRP-1A* expression in the PD brain was upregulated only during TP and CU inhibits the same

*IRP-1A* like its mammalian counterpart *IRP1* initiates iron uptake in cells, whereas high *IRP1* signalling activity is already established as a potent cause of proteinopathic NDD like AD and PD [107–110]. In the present study, *IRP-1A* expression in the PD brain is not altered during HP but upregulated by 45% (P<0.001) during TP (**Fig 5c**). CU significantly inhibited (P<0.001) the *IRP-1A* upregulation (**Fig 5c**). The observation suggests that during TP, CU intervention inhibits iron uptake by modulating *IRP-1A*, but it may not contribute to neuroprotection. Further, *IRP-1A* expression was upregulated with aging [Nearly 10 fold (P<0.001) higher in TP compared to HP] (**Fig S5c**).

#### 3.5.4. *IRP-1B* expression in PD brain was upregulated only during TP and CU partially rescues

*IRP-1B* similar to *IRP-1A* is responsible for iron intake and may also promote the progression of NDD like AD and PD [107–110]. In the present study, *IRP-1B* expression in the PD brain was not altered during HP but upregulated by 77% (P<0.001) during TP (**Fig 5d**). CU intervention partially inhibits (P<0.001) the *IRP-1B* upregulation (Fig 5D). The observation suggests that during TP, CU intervention inhibits iron uptake by modulating *IRP-1B*, but it may not contribute to neuroprotection. Further, *IRP-1B* expression was upregulated with aging [70% (P<0.001) higher in TP compared to HP] (**Fig S5d**).

## 4. Discussion

To understand the molecular basis of ALSS DAergic neuroprotective efficacy of CU we made a systematic hard effort by looking into the brain-specific modulation of molecular targets and pathways that are implicated in human DAergic neurodegeneration/neuroprotection viz., a) *dFOXO, GADD45, PUC* and proteinopathic stress under *Bsk*-signalling mediated adaptive stress response pathway; b) *IIS-dTOR* mediated anabolic signaling, mitochondrial quality control under *JNK-FOXO* and *IIS-TOR* antagonism-mediated mitochondrial dynamics; c) *CncC*, *GCLC*, *Prx 2540*, *Jafrac1* and *Prx3* of the Phase II antioxidant defense system; d) *MtnA*, *Fer1HCH, IRP-1A* and *IRP-1B* mediated brain iron metabolism. Our comprehensive work illustrates that:

### 4.1 CU-mediated ALSS neuroprotection underlies HP-specific modulation of *dFOXO, GADD45, PUC* and proteinopathic stress under *Bsk*-signalling mediated adaptive stress response pathway

*Bsk*-signalling contributes to neuronal survival, and longevity by activating antioxidant response, heat shock response, autophagy and insulin signalling antagonism, when activated in moderation [25–29], but its hyperactivation/overexpression also promotes apoptosis [25,27,111]. Further, the *Bsk-dFOXO* signalling axis is necessary for the promotion of cell survival through adaptation to stress by activating heat shock, endogenous antioxidants, growth arrest, DNA damage repair and autophagic response [28]. In the present study, *Bsk* expression in the PD brain was repressed during both the HP and TP (**Fig 1a**). It was reported that ubiquitous repression of *JNK* signalling in a *Bsk* mutant (EMS-induced) *Drosophila* model enhances susceptibility to PQ-induced stress [29]. On the other hand, it was demonstrated in *Drosophila* that neuronal enhancement of *Bsk* signalling (Heterozygous loss of *Puc* and *Bsk* overexpression) promotes resistance to PQ-induced stress, enhanced longevity, and delayed neuronal senescence [26, 29]. Concurring with the previous reports, it can be postulated in the present study that depleted *Bsk* expression is associated with PQ-mediated DAergic neuronal dysfunction and onset of PD during both the adult life stages. However, failure of CU to correct diminished *Bsk* level suggests, CU mediated HP-specific neuroprotection is not through *Bsk* modulation. Yet *Bsk* active form is the phosphorylated protein, hence further insight is needed to comment on the role of this entity in ALSS neuroprotection.

Downstream *dFOXO* expression was also diminished in PD brain of both life stages similar to upstream *Bsk*. However, it was rescued with CU intervention only during HP, but not during TP (**Fig 1b**). Tas et al, [43], reported that inhibition of *dFOXO* signalling through *RNAi* or deletion spanning in the *dFOXO* gene (Heterozygous or homozygous condition) promotes the loss of DAergic neurons and selective loss of PAM DA neuronal cluster. Therefore, onset of PD in this scenario may be associated with downregulation of *dFOXO*. *PINK1* null flies have shown significant DAergic degeneration, and mitochondrial dysfunction in the brain and thorax which was rescued with *dFOXO* and downstream gene (*SOD2*, *4e-bp*) overexpression [44]. Further, CU-analogue tetra-hydro curcumin (THC) mediated life span extension and stress resistance requires the presence of functional *dFOXO* [112]. It is hypothesized and reported that phenotypes like life span extension and DAergic neuroprotection may go hand in organisms [12, 113]. Therefore, it can be postulated from the current study that CU-mediated HP-specific neuroprotection involves *dFOXO* modulation which further down the line might modulate its target gene(s) of neuroprotection.

CU *per se* in the current study was demonstrated to enhance *Bsk* and *dFOXO* level (Compared to the control brain) only during HP (**Fig 1 a,b**). This signifies enhanced *dFOXO* signalling with CU under the physiologic condition. CU is reported as an early-acting inducer of longevity, as it enhances fly life span when fed at early stages or during HP of life through dietary means [114, 115]. Therefore, from the current study, it can be put forward that HP-specific CU-mediated modulation of *Bsk-dFOXO* during physiologic condition may contribute to longevity or better physiology. However, further insight is needed. Limitation of CU to modulate *dFOXO* during both PD and physiologic condition, *Bsk* during physiologic condition in TP may be due to diminished level of both the *Bsk* and *dFOXO* in a healthy aging brain (**Fig S1 a,b**), suggesting hindrance of their regulatory system with natural aging. It corroborates with the reports suggesting that with senescence *dFOXO* signalling activity in the muscle and neuron declines [116].

*dFOXO*-mediated *GADD45* regulation can initiate the protective DNA damage repair response by arresting cell growth and in some cases *GADD45* can also induce the pro-apoptotic signal in neuronal cells. The result demonstrated that in the PD brain of HP, *GADD45* expression is inhibited significantly (**Fig 1e**). Maitra et al, [117] (Supplementary data) reported that targeted *RNAi-*mediated knockdown of *GADD45* in DAergic neurons increases lethality of HP *Drosophila* to PQ exposure. Hence, it can be postulated that the downregulation of *GADD45* might underlie the onset of PD during HP. The current study reports that during HP diminished *GADD45* level is rescued with CU intervention (**Fig 1e**) similar to upstream *dFOXO* (**Fig 1b**).

Conditional *GADD45* overexpression (4 folds) in the fly nervous system was demonstrated to enhance stress resistance against PQ [48,50]. Therefore, CU mediated *GADD45* modulation driven by rescue of upstream *dFOXO* may contribute to life stage specific neuroprotection. However, contrasting to HP, TP PD brain demonstrated significantly increased *GADD45* expression, which was further elevated with CU intervention (**Fig 1e**). Considering the protective nature of *GADD45*, this upregulation may be a TP-specific stress compensatory response which was further boosted with CU intervention and is not correlated to upstream *dFOXO* (**Fig 1b**). But TP flies are not rescued from parkinsonian symptoms with CU intervention, suggesting the upregulation is not associated with neuroprotection. Interestingly it was reported that conditional *GADD45* overexpression (4 folds) in fly nervous system, more than 48 hrs increases mortality of flies under acute PQ (20mM) induced stress [48]. Further, 4.2-7.2 fold and 2.5-3.2 fold over expression of mammalian *GADD45γ* and *GADD45β* under nutrient deficiency was associated with neuronal apoptosis *in-vitro* [51]. Therefore, it is possible *GADD45* overexpression promotes neuroprotection in a dose dependent manner under stress condition. As observed with aging *GADD45* expression was elevated in fly brain (**Fig S1e**) that may superimpose on the upregulation of the same in TP PD brain. This may lead to negative consequence during later stages of life. Failure of CU to rescue the altered level of *GADD45* in TP PD brain highlights its limitation as a therapeutic agent against late onset NDD like PD.

HP specific modulation of *Bsk*-signalling thorough *dFOXO-GADD45* won’t be enough to promote neuroprotection if *Bsk* signalling is not balanced. The intensity of *Bsk* signalling is carefully modulated by a negative feedback loop. *Bsk* signalling-mediated regulation of *Puc* under the negative feedback loop promotes dephosphorylation of p-Bsk, preventing *Bsk* signalling overactivation and apoptosis [29,53–55]. In the current study, the result demonstrated that during HP, *Puc* expression was diminished in the PD brain (**Fig 1g**). It was reported that downregulation of *DUSP6* and *DUSP26* (A mammalian orthologue of *Drosophila Puc*), downregulation was associated with onset and progression of PD, reduced tyrosine hydroxylase, poor ATP generation, ROS production and impaired mitochondrial mobility (Rate limiting enzyme for dopamine synthesis) level [118, 119]. Hence, it can be postulated that during HP, the onset of PD, is associated with diminished *Puc* level. Further, in fly models several independent studies reported that overexpression of *Puc* suppressed irradiation and Alzheimers Disease-induced Bsk hyperactivation, alleviating morphological anomaly in wing and degenerative eye phenotype [54,55]. Therefore, it can be concluded that rescue of diminished *Puc* levels with CU in HP PD brain underlies DAergic neuroprotection. It was observed that similar to upstream *Bsk* (**Fig 1a**) CU *per se* upregulated *Puc* expression (Compared to the control brain) during HP (**Fig 1g**). It is evident that HP-specific modulation of *Puc* by CU is not limited to the PD condition, but is also present under the physiologic condition. Hence, it cannot be ruled out that CU mediated early acting longevity may also include modulation of *Puc* of the *Bsk-*signalling pathway under physiologic condition.

The unresponsiveness of *Puc* during TP suggests under PD and CU intervention suggests, *Bsk* self-regulatory axis was not utilized in any condition. Further, it was observed that aging brain had a significantly lower *Puc* level (**Fig S1g**). This insight suggests natural aging-associated hindrance of the *Puc* regulatory mechanism. During TP such natural aging-associated changes might prevent utilization of the *Bsk -Puc* axis under various conditions.

The regulation of HSP coding genes in general has been reported under the control of *Bsk* signalling and *HSP68* may be no different. *HSP68*-mediated large heat shock response is responsible for clearing toxic aggregates and promotes protein refolding in cells. The HSP coded by *HSP68* and its constitutive isoform *Hsc70* act as a chaperone and protects the cell by binding to misfolded proteins induced by oxidative stress, preventing toxic aggregation proteins [120]. In the current study, it was demonstrated that in PD brain *HSP68* expression was enhanced during both the adult life stage (**Fig 1f**). This corroborates with report of Maitra et al, [117] and considering the protective nature of the molecular player [121] suggesting neurotoxicant induces *HSP68* expression possibly as a stress compensatory response. Upon CU intervention, the altered level of *HSP68* is rescued only during HP but not during TP (**Fig 1f**). The ability of CU to inhibit *HSP68* upregulation in the PD brain suggests that CU intervention can ablate proteinopathic stress (protein misfolding and aggregation) only during HP. The current study also demonstrated that *HSP68* expression upregulated with aging (**Fig S1f**), suggesting induction of proteinopathic stress with natural aging. The onset of PD in the TP brain may further enhance cumulative proteinopathic stress. The failure of CU to impart neuroprotection in TP PD brain is due to its limitation in inhibiting proteinopathic stress. Hence, *HSP68* upregulation is also not normalized in TP PD brain.

Looking into other downstream targets of *Bsk* signalling it was found that *4e-bp* and *CAT* expression was upregulated in PD brain of both life stages (**Fig 1 d,j**). Neurotoxicant mediated upregulation of ubiquitous *4e-bp* and brain specific *CAT* in adult-young flies were previously reported [63,122]. Corroborating with these reports from the current study it can be postulated that *4e-bp* and *CAT* upregulation in PD brain are stress compensatory response. However, CU-intervention did not alter *4e-bp* level in PD brain, whereas it inhibited altered level of *CAT* in PD brain of both life phases (**Fig 1 d,j**). CU mediated inhibition of altered *CAT* level is possibly due to CU’s innate antioxidant capability. This corroborates with observation of Phom [23], suggesting although OS is sequestered during both adult life stages, it is not enough to promote neuroprotection during TP. Overall insight suggest CU-mediated HP specific neuroprotection is not through *4e-bp* and *CAT*.

Small heat shock protein *l(2)efl* and cytosolic and mitochondrial superoxide dismutase i.e. *SOD1*, *SOD2* similar differential regulatory pattern. In HP PD brain, l*(2)efl, SOD1* and *SOD2* are not altered in PD brain or with CU intervention (Fig 1 c,h,i). No alteration of *SOD1* and *SOD2* under neurotoxicant exposure in adult young fly brain was previously also reported [63]. However, in TP PD brain *l(2)efl, SOD1* and *SOD2* levels were diminished in PD brain and CU-intervention fails to rescue the same. In light of the neuroprotective and longevity extending aspect of *l(2)efl, SOD1* and *SOD2* [29,44,61], it is evident that in TP PD brain diminished level of the molecular players suggests susceptibility of TP brain to stress. Failure of CU to correct the diminished level of the players in TP PD brain highlights limitation of CU in protecting neurons at later stages of life.

### 4.2 CU-mediated ALSS neuroprotection in not through modulation of *IIS-dTOR* mediated anabolic signalling

Insulin/Insulin-like signalling (*IIS*) governs nutrient-dependent organismal growth, anabolic process, neural growth/genesis, autophagy inhibition and protein translation through activation of TOR signalling. Yet during stress or later stages of life it is more feasible curb this nergy consuming anabolic pathways in favor of energy producing pathways, thereby reducing ROS generation [30]. The current study demonstrated that *ILP*s (*Drosophila* ortholougue of insulin and isnsulin like growth factor) show variable patterns of expressional changes independent of each other. In the PD brain *ILP2* and *ILP5* were significantly downregulated (**Fig 2 a,c**), whereas *ILP3* expression was not altered during both the adult life stages (**Fig 2b**). The observation agrees with report of Karpac et al., [123] in adult young fly, which suggests such pattern of inhibition of *ILP2*,5 and unaltered level of *ILP3* is due to adaptive activation of *JNK* signalling pathway under neurotoxicant stress resulting in suppression of growth-related anabolic signalling. The current study also demonstrates that *ILP* downstream insulin receptor substrate *Chico* level was upregulated in the PD brain of both adult life stages (**Fig 2d**). A high sugar diet is reported to promote Type 2 Diabetes Mellitus in adult young (15 days old) *Drosophila*, resulting in ubiquitous upregulation *Chico* and induction of motor deficit [124]. Therefore, it is possible upregulated *Chico* in the PD brain may lead to some form of metabolic consequences contributing to onset and progression of PD. CU intervention inhibited all the *ILP*s level and corrected alter *Chico* level in PD brain of both life stages (**Fig 2 a-d**). Aging, metabolic response/disorder and neurodegeneration go hand in hand and components of *IIS* actively regulate the process [31]. Ubiquitous inhibition of *ILP*s and *Chico* expression through *RNAi* mediated knock down and gene knock out respectively promotes extended lifespan, and stress resistance in fly models [69,125]. However, despite the positive implications of *IIS* i.e., *ILP*s and *Chico* inhibition TP flies are not rescued with CU intervention. It is possible that inhibition mediated longevity extension and DAergic neuroprotection do not go hand in hand, therefore CU mediated HP-specific neuroprotection may not be through modulation of *IIS*.

To further understand the role of the anabolic signalling in ALSS neuroprotection we investigated the differential modulation of *dTOR* (Fly orthologue of mTOR) and *RS6K1*. The current study demonstrates that *dTOR* expression in PD brain was significantly enhanced during HP, whereas expression of the same was not altered during TP (**Fig 2e**). In the adult young (2-3 months old) mice model, it was demonstrated that PQ exposure results in the onset of PD with an enhanced mTOR translate level [126]. The current observation in the HP PD brain corroborates with the finding of Wills et al, [126] suggesting higher *dTOR* transcript level may in fact be associated with the onset of PD. However, during TP such upregulation in PD is not observed, but the possibility of higher dTOR kinase activity cannot be ruled out as *Chico* of the upstream *IIS* signalling is upregulated in the TP PD brain (**Fig 2d**), suggesting enhanced signalling of *IIS*. CU intervention during HP normalizes *dTOR* altered level, whereas inhibits the same during TP (**Fig 2e**). Dietary intervention of CU is shown to promote life span extension in *Drosophila* through *dTOR* inhibition [127]. Further, ubiquitous overexpression of a dominant negative form of *dTOR* increases the *Drosophila* life span [70]. However, flies of TP are not rescued with CU intervention from PD symptoms, therefore it is possible DAergic neuroprotection by CU does not underlie modulation of *dTOR* similar to upstream *IIS*.

It must be noted however that the active form of *IIS-dTOR* pathway is phosphorylated Chico, dTOR and RS6K1. Hence, further insight is needed to comment on the role of this pathway in CU-mediated ALSS neuroprotection.

### 4.3 CU-mediated ALSS neuroprotection in through rescue of mitochondrial quality control under *JNK-FOXO* and *IIS-TOR* antagonism-mediated mitochondrial dynamics

Anabolic processes require higher energy consumption; hence mito-biogenesis must be increased. Insights suggest that *mTOR/dTOR* signalling of the *IIS-mTOR/dTOR* pathway can also promote gene expression and translation of transcription factor coding *Ewg* (*NRF1* as mammalian orthologue), which down the line regulates the expression of *TFAM*. *TFAM* promotes the replication and transcription of mtDNA necessary for mito- biogenesis [71,72]. The current study demonstrated that in the PD brain of both adult life stages, *Ewg* expression is downregulated and CU intervention fails to rescue (**Fig 3a**). Neurotoxicant like MPTP and MPP^+^ (Chemically very similar to PQ) is reported to inhibit *NRF1* expression, and translate level in neuronal cell culture and in mice brain resulting in PD condition [128, 129]. The current observation in *Drosophila* PD brain corroborates the insights from Wang et al, and Piao et al, [128, 129] suggesting that diminished *Ewg* is associated to onset of PD. As *Ewg* also expresses electron transport chain (ETC) subunits [130], diminished *Ewg* may also give cues to deficiency in ATP production. Failure of CU to rescue deficient *Ewg* level during both life stages suggest, CU-mediated ALSS neuroprotection is not through *Ewg*.

On the other hand, to obtain further insight into the mito-biogenesis under *IIS-dTOR* signalling, expression analysis of *TFAM* was also performed. The current study demonstrated that during both the adult life stages *TFAM* expression is not altered in the PD brain, yet CU intervention inhibited *TFAM* (**Fig 3b**). CU mediated inhibition of *TFAM* is like inhibition of upstream anabolic *IIS-dTOR* (**Fig 2 a-e**). In a mice model of AD, CU intervention had contrasting action on *TFAM* expression. It was demonstrated that depending on the mode of toxicity with two different variants of ApoE–4, CU either upregulated *TFAM* or downregulated it to promote neuroprotection and uplift mitochondrial health [131]. Hence, it is highly likely that CU-mediated rescue of mitochondrial biology through *TFAM* modulation is highly adaptive to the nature of toxicity and degenerative condition. Further, when mitochondrial biogenesis-specific signalling is enhanced, it leads to neural degeneration in the fly model. *RNAi*-mediated *TFAM* inhibition is demonstrated to be neuroprotective in such cases of aberrant mito- biogenesis [73,132]. In the current study, enhanced it is evident that during time of stress CU intervention also suppresses mito-biogenesis along with upstream *IIS-dTOR* anabolic signalling. This might have a protective aspect as it may prevent propagation of faulty mitochondrial DNA, but TP flies are not rescued with CU intervention. Therefore, CU-mediated ALSS neuroprotection is not through modulation of *TFAM*.

With the insight on *IIS-dTOR* anabolic signalling mediated mito-biogenesis in ALSS neuroprotection, We further investigated the mitochondrial quality control antagonistic to mito-biogenesis and is controlled by *Bsk-dFOXO* pathway [33,34,74–76]. Like upstream *dFOXO* (**Fig 1b**), diminished level of *Mfn2* in PD brain was rescued with CU- intervention only during HP but not during TP (**Fig 3c**). In young fly model and mice model neurotoxicant exposure resulted in diminished *Mfn2* level (Transcript and translate) resulting in oxidative stress, motor deficit, mitochondrial fragmentation in thoraces and brain respectively [74,75]. PD patients also show a trend toward downregulated Mfn2 translate level compared to age-matched controls [75]. In an *in-vitro* neuronal cell model of AD, CU is reported to enhance *Mfn2* expression and promotes protection against amyloid beta toxicity [133]. Further, *Mfn2* overexpression in mice model is reported to ablate MPTP-mediated neurodegeneration in the *substantia nigra* of mice model [134]. Also, *FOXO*-mediated mitochondrial homeostasis is documented for neuronal and non-neuronal tissues [76,135]. Therefore, considering the evidences, insight from the current study suggests that CU-mediated differential modulation of *Mfn2* driven by *dFOXO* underlies the ALSS neuroprotection. Like *dFOXO* (**Fig 1b**) CU *per se* is also shown to enhance *Mfn2* level only during HP (**Fig 3c**). Therefore, it is possible CU-mediated early life acting longevity enhancement, also underlies modulation of *Bsk-dFOXO* axis mediated *Mfn2* modulation under the physiologic condition.

To further understand mitochondrial respiratory capacity differential modulation *ATP SynD* was analysed which codes for a subunit-D of complex -V of respiratory chain. Onset of PD does not alter *ATP SynD* level during both the life stages (**Fig 3d**). Neurotoxicant PQ is not known for selectively inhibiting mitochondrial complexes [136], therefore the current observation corroborates the reports. CU intervention is shown to enhance *ATP SynD* level only during HP (**Fig 3d**). CU itself is an active mediator of ATP biogenesis [137], but enhancement of *ATP SynD* during in HP PD brain with CU intervention suggests that the respiratory capacity of neurons are elevated possibly due to overall betterment of mitochondrial health.

### 4.4 CU-mediated ALSS neuroprotection in through HP specific modulation of *CncC*, *GCLC*, *Prx 2540*, *Jafrac1* and *Prx3* of the Phase II antioxidant defence system (ADS)

*CncC* (*Drosophila* orthologue of mammalian *Nrf2*) is a mediator of the phase II ADS which is responsible for the neutralization of the xenobiotics, toxins, and induced peroxide stress [78–80]. Further, some of the ADS components can be regulated by Bsk- *dFOXO* signalling [29,49,93]. In the present study, *CncC* expression in the PD brain was inhibited during HP and TP, whereas rescued with CU intervention only during HP (**Fig 4a**). It was reported in *in-vitro* neuronal cells that exposure to sub-lethal PQ leads to depletion of *Nrf2* which leads to impaired redox balance [81]. Further, *CncC* overexpression in the DAergic neuron of *Drosophila* ablates α-synuclein mediated toxicity, and neuronal death and enhances fly survival under harsh conditions [78]. Hence, *CncC* is highly necessary for optimal neuronal health and downregulation of the same is associated with the onset of PD. CU is a known modulator of *Nrf2* in various neuronal disorders. CU and/or anti-cholinesterase combined formulation rescues diminished *CncC* expression and improves memory impairment in a young *Drosophila* model [138]. In a young mice model of focal ischemia and traumatic brain injury, CU intervention is demonstrated to alleviate neurodegenerative phenotype by rescuing diminished *Nrf2* transcript level resulting in elevated level of downstream antioxidant gene expression [139]. Therefore, CU intervention promotes neuroprotection by rescuing the diminished level of *CncC* only during HP. Resuscitation of the *CncC* may contribute to the elevation of phase II ADS resulting in better neuronal defence during health phase but similar is not possible during TP.

In the present study, *GCLC* expression was downregulated in the PD brain during HP, which was rescued upon CU intervention. But during TP there was a significant upregulation of *GCLC* in the PD brain which could not be altered with CU intervention (**Fig 4b**). *GCLC* expression is highly responsive to *Nrf2/CncC* signalling. Like *CncC* sublethal PQ exposure decreases *GCLC* level, leading to impaired redox balance [81]. Further, increased GCLC translate level due to *CncC* inducer treatment or DAergic *CncC* overexpression ablates α-synuclein mediated toxicity, and neuronal death in *Drosophila* [78]. Therefore, it is evident that CU mediated rescue of diminished *GCLC* driven by *CncC* underlies HP specific neuroprotection. However, *GCLC* upregulation in TP PD brain may be stress compensatory response, failure to rescue the altered level of the same by CU intervention suggest inability of CU to curb DAergic stress during later stages of life. Similarly, *Jafrac1* expression in HP PD brain was downregulated which was rescued with CU intervention (**Fig 4f**). Lee et al, [93] also demonstrated that *RNAi-*mediated neuronal knockdown of *Jafrac1* enhanced PQ-induced lethality in flies, whereas neuronal overexpression of the same reduced PQ-induced lethality. In DA neuronal cell culture and mice brains, it was reported that overexpression of *Prx2* (A mammalian orthologue of *Jafrac1*) sequesters 6-OHDA-induced oxidative stress and neuronal death [140]. Further, *Jafrac1* is also modulated my *dFOXO* [93], like *CncC*. Therefore, CU mediated HP-specific neuroprotection may underlie rescue of diminished *Jafrac1* possibly driven by upstream *dFOXO* and/or *CncC*. In the PD brain of TP, expression of *Jafrac1* is enhanced, which is further enhanced with CU intervention (**Fig 4f**). Considering the protective role of *Jafrac1* it is possible that such upregulation in TP PD brain is an adult life stage-specific adaptive response to stress which is further enhanced by CU and is not driven by *dFOXO* and/or *CncC*. However, this enhanced *Jafrac1* level may not protect from neurodegeneration during TP. Rescue of *GCLC* and *Jafrac1* in HP suggest that reduced-GSH production is rescued and peroxide neutralization under the command of phase II mediator *CncC* and/or adaptive stress response mediator *dFOXO*.

Interestingly the cross talk between adaptive stress response mediator *dFOXO* and phase II mediator *CncC* also extends to mitochondrial peroxiredoxins. *Prx5* and *Prx3* are two mitochondrial peroxiredoxins, where known modulator of *Prx5* is *CncC*, *Prx3* can also be modulated by *dFOXO* other than *CncC*. *Prx5* product is ubiquitous to both mitochondrial and cytosol [36]. In the present study, *Prx5* expression in the PD brain was inhibited during HP and TP which was rescued with CU intervention (**Fig 4h**). It was reported that *Prx5* null flies show shortened life span by itself or with stress [97]. Further, global overexpression of the *Prx5* increased resistance to oxidative stress and enhanced fly life span by 30% in normal conditions [97]. Therefore, it can be the inhibition of *Prx5* during both the adult life stages is associated with the onset of PD, which was rescued with CU intervention. Yet, TP flies are not rescued with CU intervention, hence the ALSS neuroprotection is not through *Prx5* modulation.

*Prx3* on the other hand codes for only mitochondria specific antioxidant [36]. Diminished level of the same under neurotoxicant insult leads to onset of PD in-vitro and neuronal over expression of the same in fly model enhances stress resistance particularly in middle age [96,141]. The current observation suggests like upstream *dFOXO* and *CncC*, diminished *Prx3* level was rescued with CU intervention only during HP but not during transition phase (**Fig 4i**). Therefore, insight suggests that diminished *Prx3* level may lead to poor mitochondrial antioxidant capability in PD brain, which was corrected with during HP with CU intervention driven by *dFOXO* and/or *CncC*.

In the population of good peroxiredoxins under the phase II ADS one acts as a double-edged sword. *Prx 2540* or *Prx 2540-1,2* other than having antioxidant capability has potent inflammatory capability [36]. *Prx 2540* in *Drosophila* exist as two identical copies *viz. Prx 2540-1* and *Prx 2540-2* (*Prx 2540-1,2*). In the present study, *Prx 2540-1,2* expression was upregulated in the PD brain, whereas CU intervention normalized the altered expression of the same only during HP (**Fig 4e**). Due to its reported antioxidant capability [91], it can be postulated that upregulation of the same in PD brain is a compensatory response to stress. Upon inhibition of stress in HP the altered *Prx 2540-1,2* level is rescued. But, it was also reported that *PRDX6* (Mammalian orthologue of *Prx 2540*) transgenic mice show more DAergic neuronal loss, reduced tyrosine hydroxylase level and persistant inflammation upon MPTP exposure [142]. Therefore, in the current study, it is possible that in PD brain *Prx 2540-1,2* upregulation may underlie DAergic neuronal degeneration and reduced tyrosine hydroxylase level as previously reported [143]. Owing to CU’s anti-inflammatory nature, intervention of the same inhibits *Prx 2540-1,2* during HP and protects the neuron, but same is not possible in TP PD brain. Therefore, CU mediated ALSS neuroprotection underlies HP specific modulation of *Prx 2540-1,2*.

Similarly, *GCLM* was upregulated in the PD brain during HP and TP, yet the altered level was rescued only during HP (**Fig 4c**). *CncC* downstream *GCLM* stabilizes *GCLC*- mediated GSH production [82]. *GCLM* is shown to promote neuroprotection and stress resistance [83, 144]. Therefore, the observed upregulation of the same in PD brain may be a stress compensatory response. CU could rescue the altered level of *GCLM* during HP but not during TP, suggesting CU can ablate some form of stress combated by *GCLM* only during HP, resulting in correction of the altered *GCLM* level under stress.

*GSTD1* of the phase II ADS is also under the control of *Bsk* signalling pathway [29]. It neutralizes xenobiotics and herbicides and shown to be upregulated in their presence in fly model [62,85,88]. Our observation corroborates the previous reports as in PD brain of both life stages *GSTD1* level is enhanced, suggesting it is a general stress responder. CU intervention in PD brain does not alter the *GSTD1* level, eluding that CU mediated ALSS neuroprotection is not through *GSTD1*.

### 4.5 CU’s limitation in TP may underlie its failure to inhibit high iron intake signals in PD brain

It was reported that metal (copper, heavy metal and iron) chelator protein-coding genes *MtnA* and *Fer1HCH* may respond to *Bsk* signalling in the *Drosophila* [29]. In the present study, *MtnA* and *Fer1HCH* expression in the PD brain was upregulated, while CU intervention could partially rescue the altered level during both the adult life stages (**Fig 5 a,b**). *Mt-II* (Vertebrate orthologue of fly *MtnA*) has previously been reported to be upregulated under neurotoxicant stress, traumatic injury in brain, which corroborates with our observation [100, 102]. *MtnA* are known for their role in heavy metal detoxification, anti-inflammatory and antioxidant role, that promotes neuroprotection against proteinopathic NDD like AD [145, 146]. Similarly, *Fer1HCH* is necessary for maintaining free iron chelation, preventing ROS generation in nervous system. Further, in *Drosophila Fer1HCH* is known to prevent axonal degeneration of axons and protect nervous system against ferroptosis [103, 104]. Therefore, upregulated *MtnA* and *Fer1HCH* in PD brain may be a stress compensatory response. Similarly high *Fer1HCH* expression is also observed in aging brain in *Drosophila* (**Fig S5b**), suggesting mechanism of aging and PD onset both leads to iron accumulation stress. CU intervention during both the adult life stages partially rescue the upregulated *MtnA* and *Fer1HCH* levels, suggesting that CU intervention in the PD brain can somewhat prevent metal imbalance-associated stress. This action by CU may be due to its inherent chemistry, that enables it to perform as a metal chelating molecule [147]. However, this may not be enough to neuroprotection during later stages of life as observation suggests.

Therefore, to further explore the reason behind inefficacy of CU we investigated the iron intake components of this metal homeostasis pathway. The enhanced *IRP* signalling results in the inhibition of *Fer1HCH* mediating iron chelation and simultaneously enhancing iron intake through activation of *Tfr1* signalling [107]. In the present study, *IRP-1A* and *IRP-1B* expression in the PD brain was unaltered during HP, but expressions of the same were upregulated during TP. However, during TP CU intervention fully inhibited *IRP-1A* upregulation but *IRP-1B* upregulation was only partially rescued (**Fig 5 c,d**). In PD and AD iron intake mediated by high IRP-IRE binding activity, has been a leading cause of the NDD [108,110]. Therefore, it is evident from the present study that in the PD brain upregulation of *IRP-1A* and *IRP-1B* during TP may contribute to the enhanced detriment in neurons. Further, natural aging also promoted extremely high *IRP- 1A* and significantly high *IRP-1B* expression in fly brains (**Fig S5 c,d**). Thereby, suggesting cumulative iron intake signalling is enhanced upon PD onset during TP. CU intervention although inhibits *IRP-1A* upregulation, it fails to completely rescue *IRP-1B* upregulation. Therefore, even though PD-associated *IRP-1A* upregulation is inhibited fully, the partial rescue of *IRP-1B* is not enough to curb the cumulative high *IRP* signalling intensity during TP. Hence, high iron intake and oxidative stress may still prevail during TP, highlighting one of the possible reasons for CU inefficacy during later adult life stages.

## 5. Conclusion

Insights from the present study illustrate that possible players of CU-mediated HP-specific neuroprotection belong to stress-responsive *Bsk* signalling pathway, mitochondrial dynamics pathway and phase II ADS pathway. Further, these networks may have a potent cross-talk as different networks regulate similar downstream targets and/or their action converges on the mitochondria (**Fig 6**). The molecular players of CU mediated HP specific neuroprotection are (**Fig 6**)

- *Bsk-dFOXO* stress response pathway: *dFOXO, GADD45, Puc*
- Mitochondrial dynamics: *Mfn2*
- Phase II ADS pathway: *CncC, GCLC, Prx 2540 -1,2, Jafrac1, Prx3*

**Figure 6:**
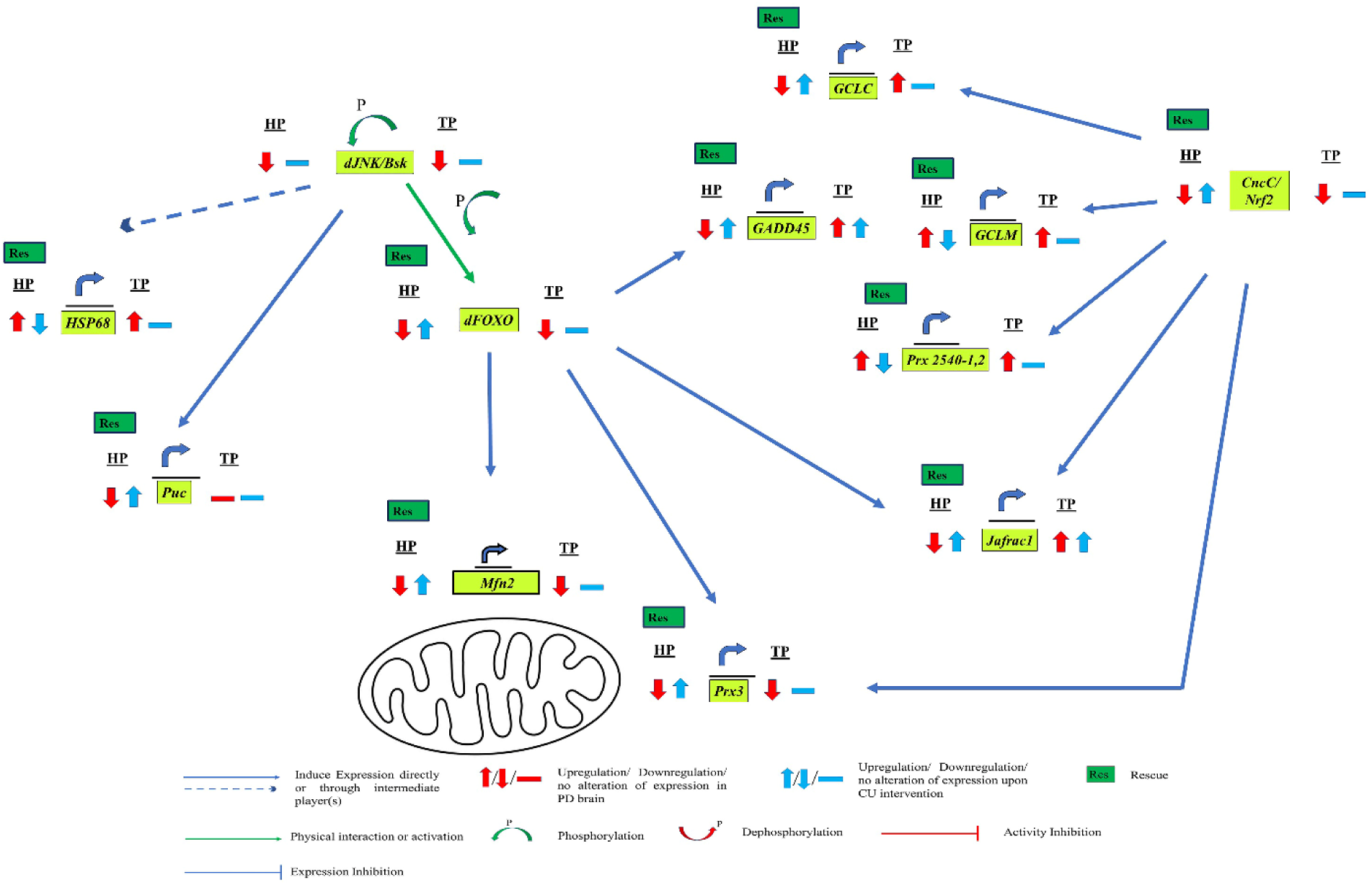
Cartoon depicts the involvement of molecular players in the HP-specific DAergic neuroprotective efficacy of CU. Insights reveal that CU-mediated ALSS DAergic neuroprotection underlies differential modulation of the molecular players belonging to the *Bsk* signalling pathway, mitochondrial dynamic pathway, and phase II ADS pathway.

Failure to rescue the altered level of these players in TP PD brain suggests CU’s inefficacy during later stages of life. Further, CU-mediated inhibition of proteinopathic stress and some stress countered by Phase II ADS leads to the correction of altered levels of *HSP68* and *GCLM* during HP only. Since, CU mediated HP specific neuroprotection promotes mitochondrial quality control in a *dFOXO-Mfn2* dependent pathway and mitochondrial anti-oxidant capability in a *CncC* and/or *dFOXO* – *Prx3* dependent pathway, it is concluded that mitochondrial health modulation may underlie life stage-specific neuroprotection.

We would further like to emphasize that since CU’s inception as a phytomedicine/therapeutic molecule and despite its shortcomings as a therapeutic agent in PD [10], still to these days CU has been proposed as a phytomedicine and one of the potential alternatives to currently available drugs when at least NDDs are concerned [148]. Perhaps a relook is necessary on how this potent nutraceutical be utilized to counter a late-onset diseases. Our study highlights that it is the natural aging associated molecular changes in neuro-biological pathways that renders therapeutic efficacy of CU as null in late-onset NDD such as PD. On the other hand, not therapeutic dosage, but dietary intervention of CU is a life span extender when fed at early life stages [115, 127], that is achieved through regulation of various molecular players. Perhaps if the molecular players can be sustained at a healthy regulatory level by dietary intervention of the nutraceutical, then the onset of the late-onset PD be prevented. Our lab is currently working in that direction. Further we are also working to sustain the observed brain-specific molecular players/pathways that confer DAergic neuroprotection during HP, in TP by adhering to different life course mediated feeding regimes. This understanding will help to modify existing therpapeutic strategies and also to develop novel therapeutic methods for late-onset NDDs such as PD.

## Acknowledgements

Author Contributions:

Conceptualization:SCY Data curation:SCY, AD Formal analysis:AD Funding acquisition:SCY Investigation:AD, SCY Methodology:SCY Project administration:SCY Resources:SCY Software:AD, SCY Supervision:SCY Validation: AD, SCY Visualization: AD, SCY Writing - original draft:AD Writing - review & editing:SCY

## Data Availability

The authors declare that the data supporting the findings of this study are available within the paper and its Supplementary Information files. Should any raw data files be needed in another format they are available from the corresponding author upon reasonable request.

## Declarations

**Ethics Approval and Consent to Participate:** Not applicable.

**Consent for Publication:** Not applicable.

**Competing Interests:** The authors declare no competing interests

## Abbreviation

CU: Curcumin DAergic: Dopaminergic
HP: Health Phase *SNpc: Substantia Nigra pars compacta*
TP: Transition Phase DA: Dopamine
PD: Parkinson’s Disease
L-DOPA: Levodopa
NDD: Neurodegenerative diseases
ALSS: Adult life stage specific

---

**Figure S1:**
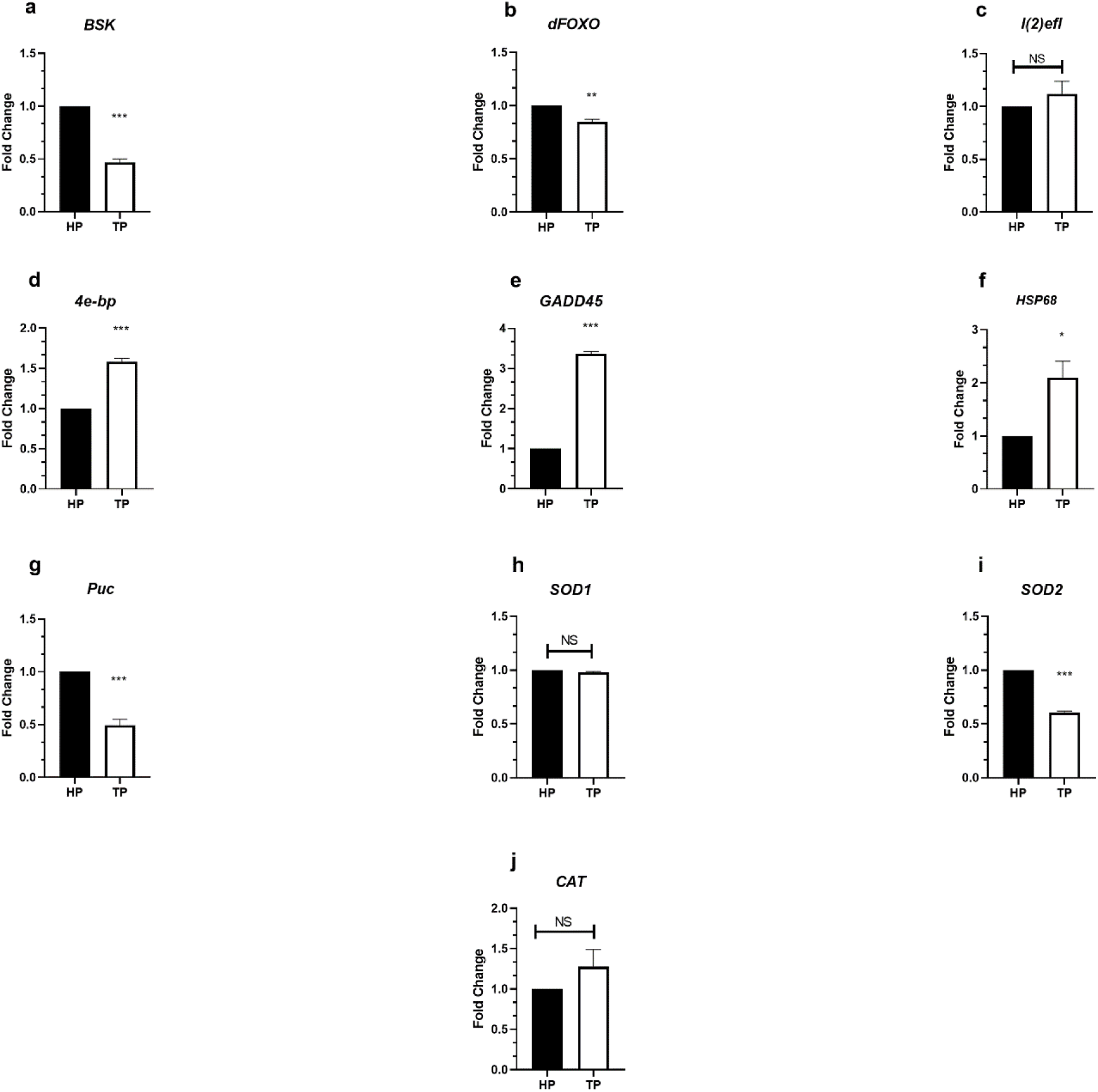
Brain-specific aging associated changes in expression of molecular players involved in *Bsk* signalling pathway. Natural aging in brain inhibits *Bsk-dFOXO* stress response axis, as expression level of *Bsk* and *dFOXO* was inhibited in TP brain as compared to that of HP (a,b). However, *dFOXO* downstream *l(2)efl* expression was unaltered, whereas *4e-bp* and *GADD45* expression were upregulated in TP brain as compared to HP (c,d,e). Similarly *Bsk* downstream *HSP68* expression was also enhanced, whereas *Bsk* downstream *Puc* expression was inhibited in TP brain as compared to HP (f,g). *dFOXO* downstream antioxidant gene *SOD1* and *CAT* remain unaltered, whereas *SOD2* expression was inhibited in TP brain as compared to HP (h,i,j). Significance was drawn by analysing the data of minimum three replicates with unpaired t-Test. (*p<0.05; **p<0.01; ***p<0.001; NS: Not significant)

**Figure S2:**
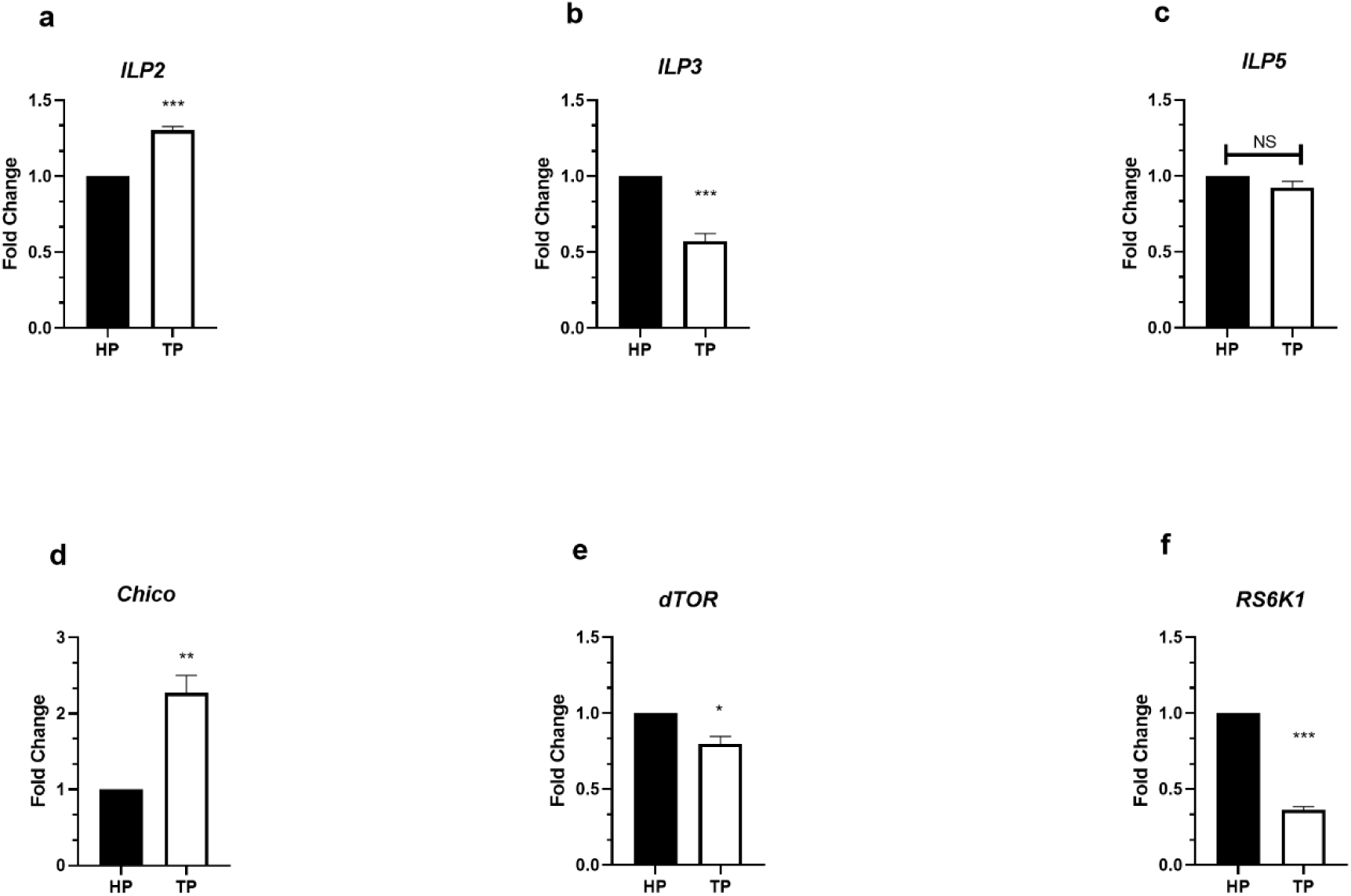
Brain-specific aging associated changes in expression of molecular players involved in *IIS- dTOR* signalling pathway. Natural aging in the brain may enhance cumulative *IIS* signalling as expression level *ILP2* was enhanced, whereas *ILP3* was downregulated and *ILP5* remained unaltered in TP brain as compared to that of HP (a,b,c). Similarly, *ILP* downstream *Chico* level was also upregulated (d). However, unlike *IIS,* the downstream *dTOR-RS6K1* signalling cascade may be repressed with natural aging as *dTOR* and RS6K1 level was inhibited in the TP brain as compared to that of HP (e,f). Significance was drawn by analysing the data of minimum three replicates with unpaired t-Test. (*p<0.05; **p<0.01; ***p<0.001; NS: Not significant)

**Figure S3:**
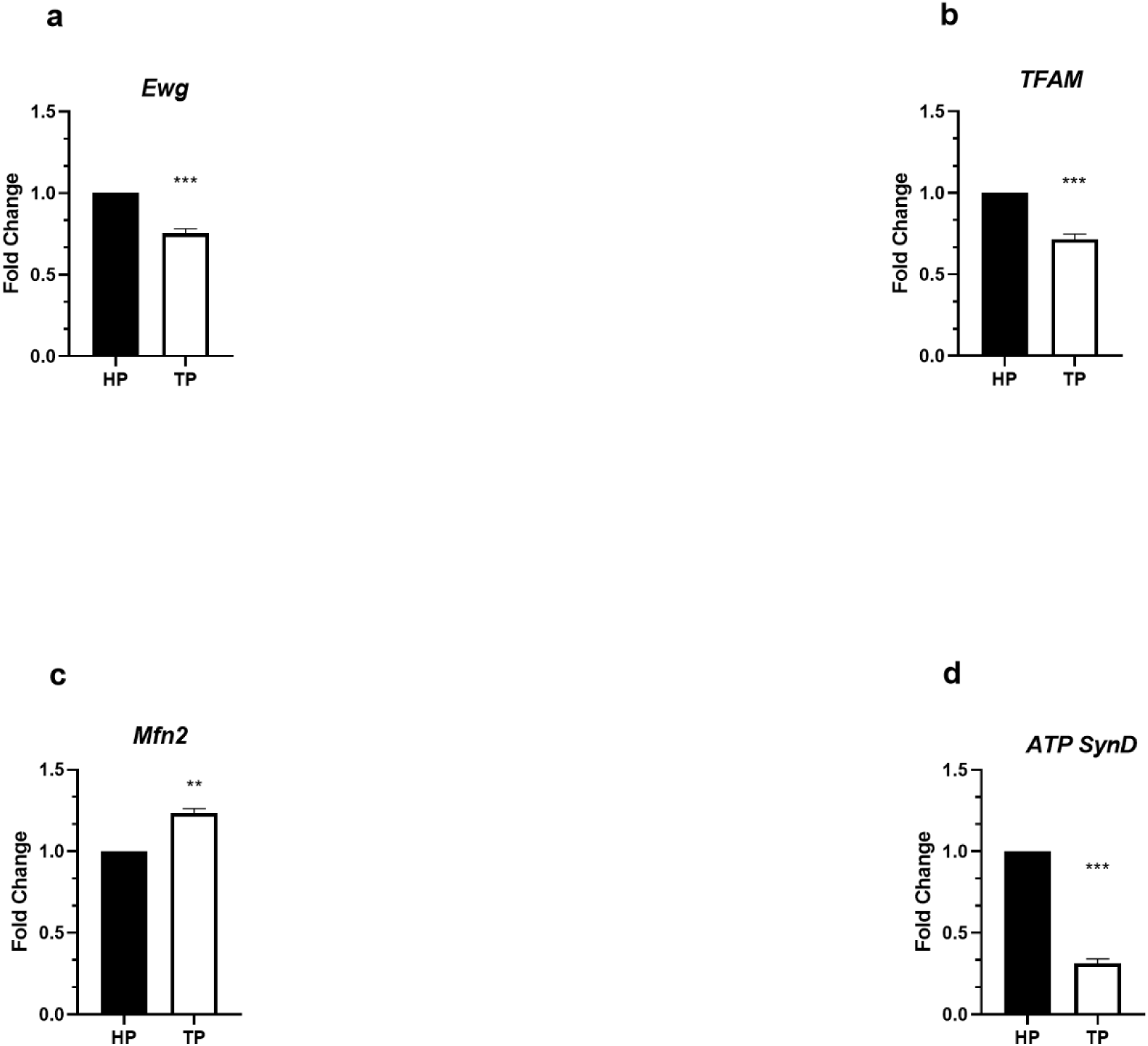
Brain-specific aging associated changes in expression of molecular players involved in mitochondrial dynamics. Natural aging in brain inhibits mito-biogenesis capacity as *Ewg* and *TFAM* expression level were inhibited in the TP brain as compared to that of HP (a,b). However, mito-quality control is enhanced with natural aging as *Mfn2* level was upregulated in the TP brain as compared to that of HP (c). Natural aging diminishes respiratory capacity in the brain as *ATP SynD* was diminished in the TP brain as compared to that of HP (d). Significance was drawn by analysing the data of minimum three replicates with unpaired t-Test. (*p<0.05; **p<0.01; ***p<0.001; NS: Not significant)

**Figure S4:**
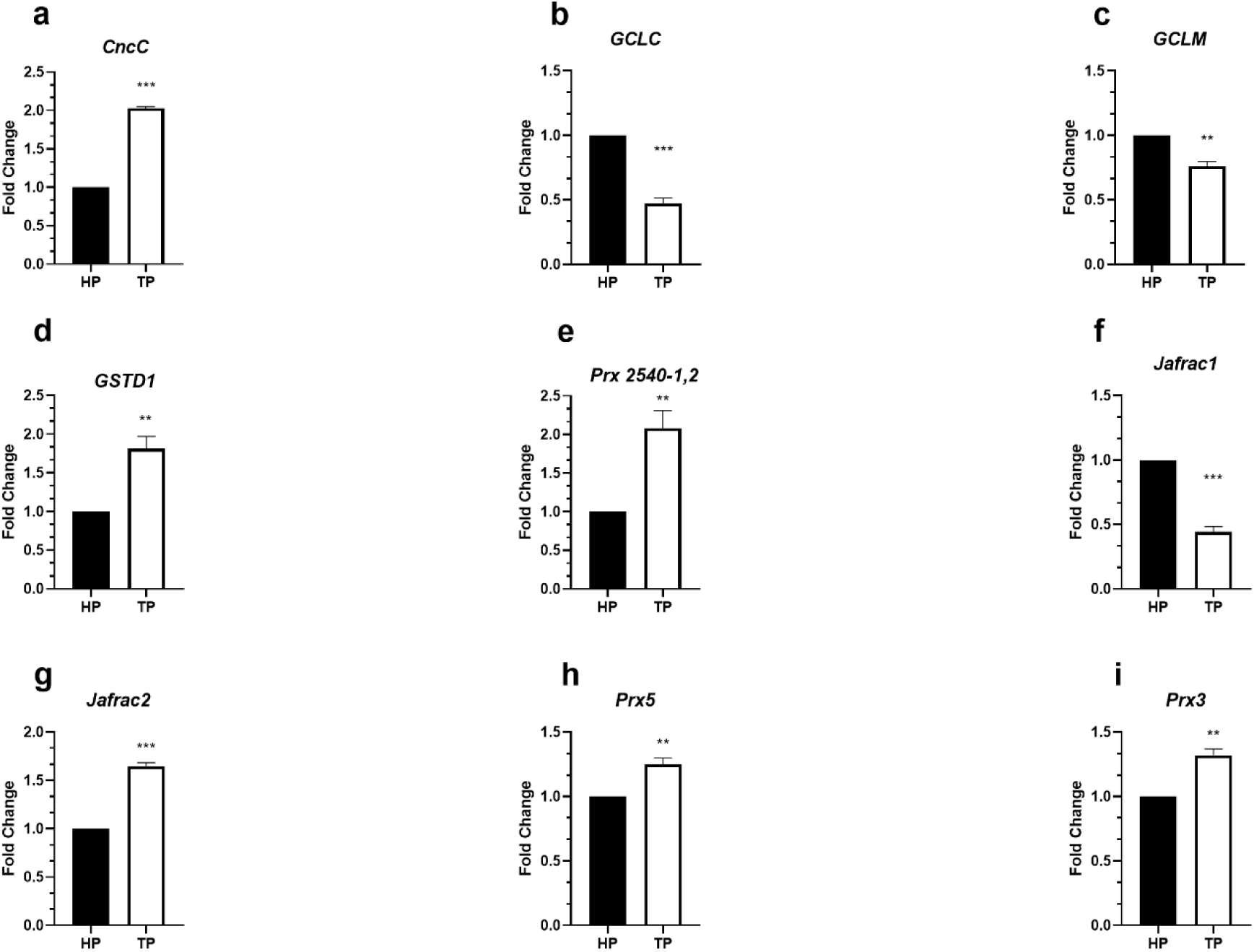
Brain-specific aging associated changes in expression of molecular players involved in phase II ADS. Natural aging in the brain upregulates phase II ADS mediator *CncC* (a). However, *CncC* downstream *GCLC* and *GCLM* were downregulated in the TP brain, suggesting a decline in GSH synthesis with aging (b,c). Further, *GSTD1*, *Prx 2540-1,2*, *Jafrac2*, *Prx5* and *Prx3* were upregulated in TP brain, whereas *Jafrac1* was downregulated (d,e,f,g,h,i). Significance was drawn by analysing the data of minimum three replicates with unpaired t-Test. (*p<0.05; **p<0.01; ***p<0.001; NS: Not significant)

**Figure S5:**
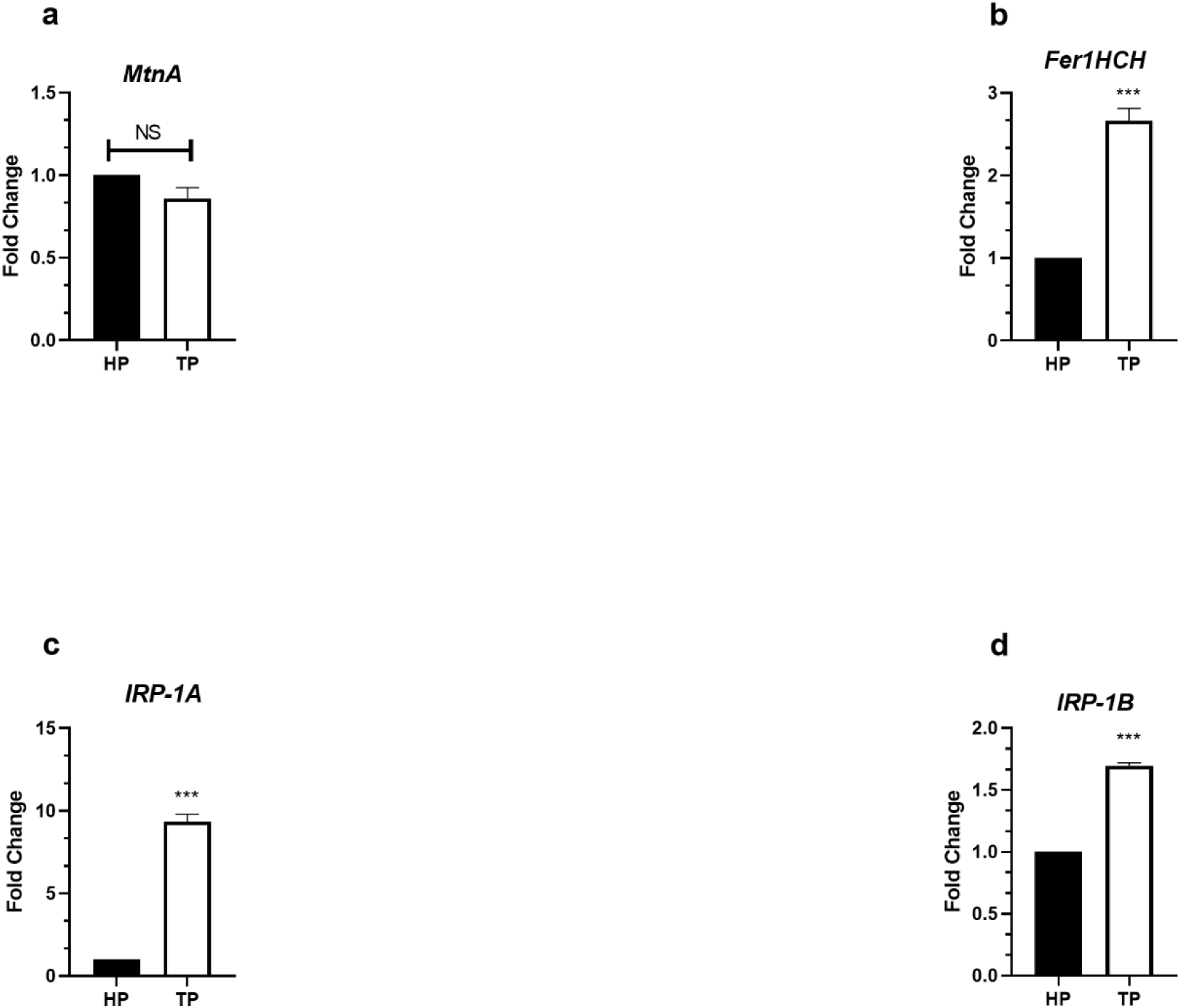
Brain-specific aging associated changes in expression of molecular players involved in metal homeostasis. Natural aging in the brain did not alter *MtnA* expression (a). But with aging iron accumulation in the brain may be enhanced and in response to that *Fer1HCH* was upregulated in TP brain (b). Similarly in the TP brain iron uptake was also enhanced as *IRP-1A* and *IRP-1B* expression was upregulated (c,d). Significance was drawn by analysing the data of minimum three replicates with unpaired t-Test. (*p<0.05; **p<0.01; ***p<0.001; NS: Not significant)

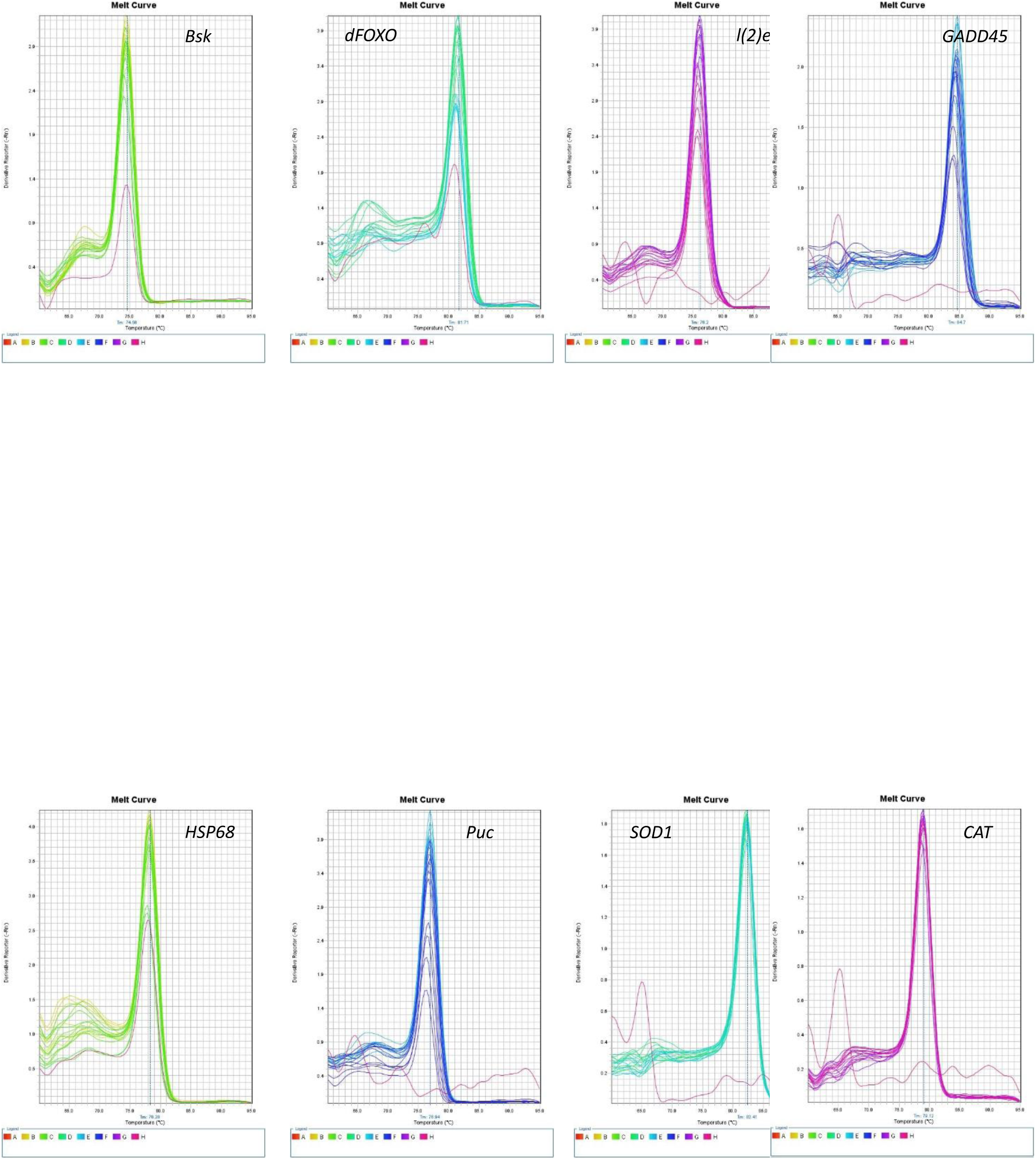
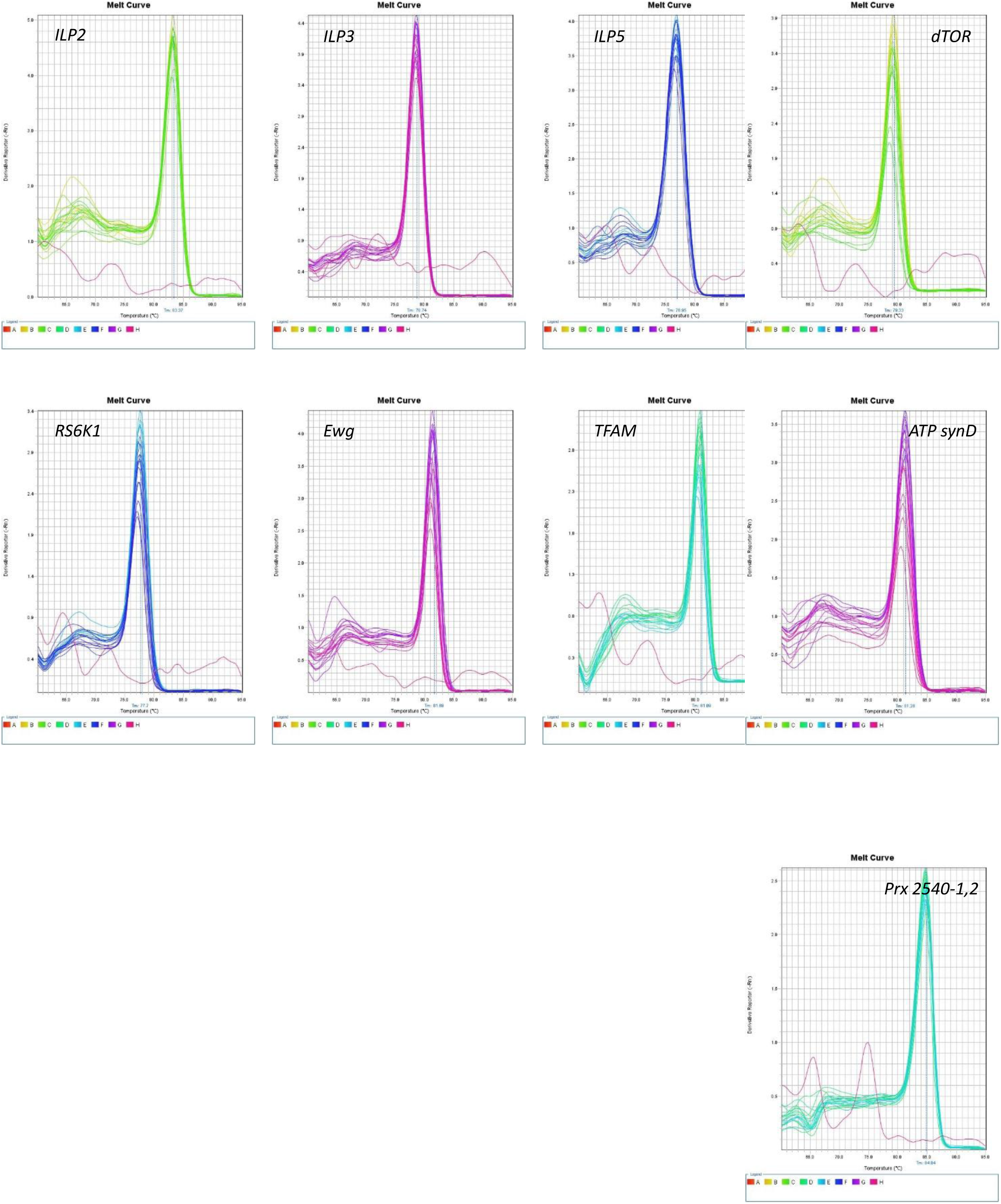
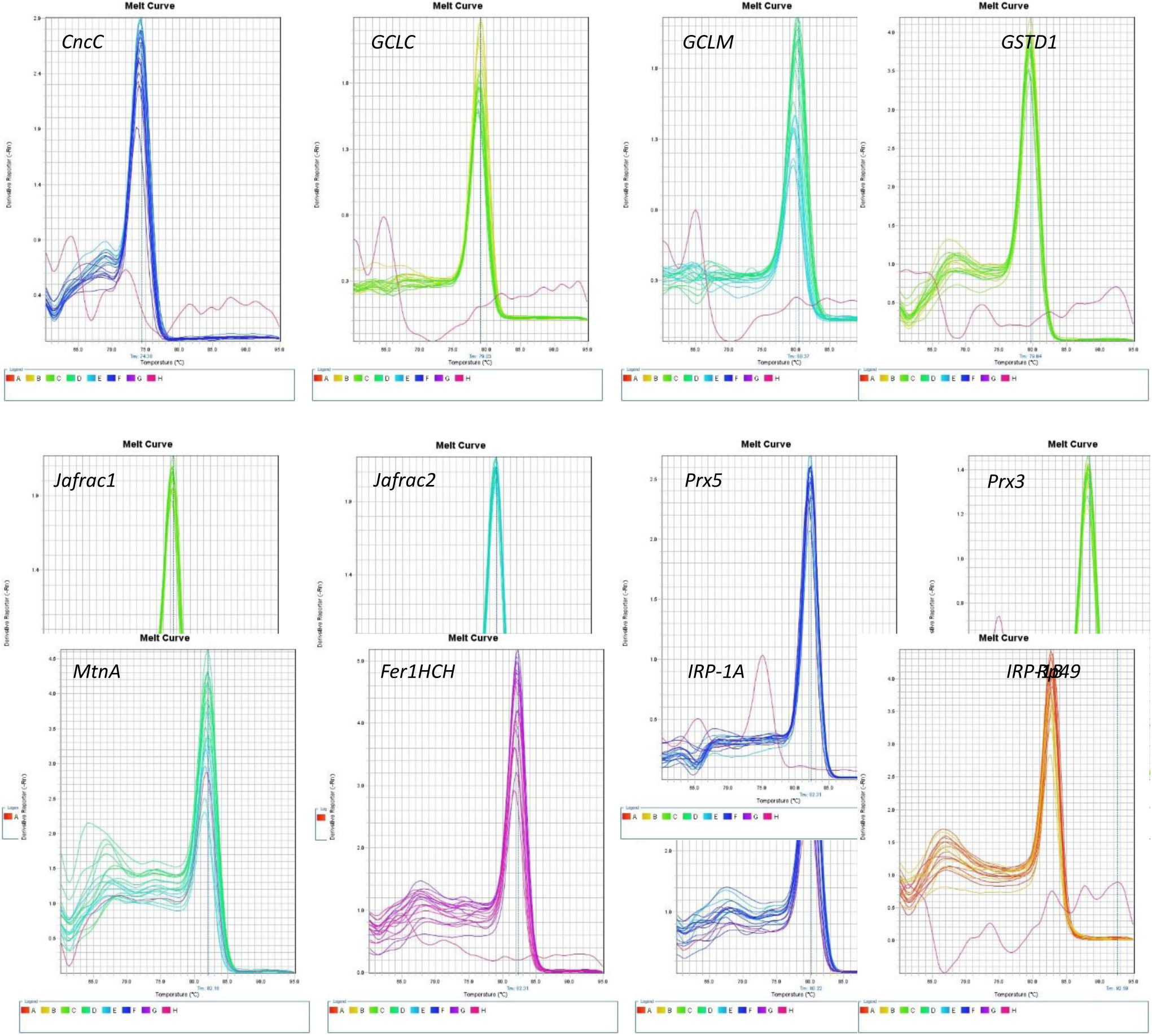

