## Supplementary data for "Differential regulation of brain-specific molecular pathways is the reason for curcumin’s adult life-phase specific DAergic neuroprotection: Insights from ALSS Drosophila model of Parkinson’s disease"

### Supplementary Figures

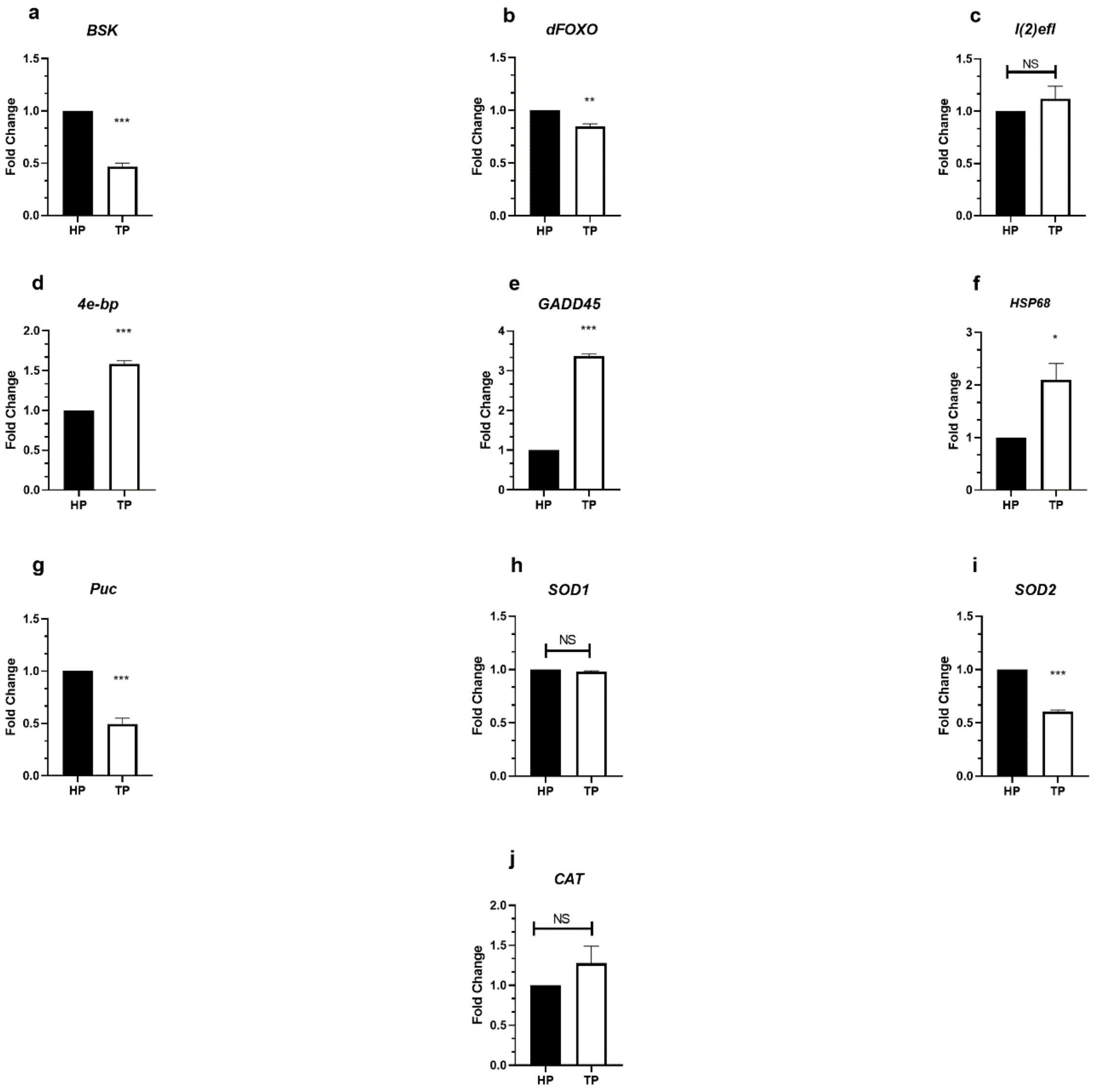


**Figure S1**: Brain-specific aging associated changes in expression of molecular players involved in *Bsk* signalling pathway. Natural aging in brain inhibits *Bsk-dFOXO* stress response axis, as expression level of *Bsk* and *dFOXO* was inhibited in TP brain as compared to that of HP (a,b). However, *dFOXO* downstream *l(2)efl* expression was unaltered, whereas *4e-bp* and *GADD45* expression were upregulated in TP brain as compared to HP (c,d,e). Similarly *Bsk* downstream *HSP68* expression was also enhanced, whereas *Bsk* downstream *Puc* expression was inhibited in TP brain as compared to HP (f,g). *dFOXO* downstream antioxidant gene *SOD1* and *CAT* remain unaltered, whereas *SOD2* expression was inhibited in TP brain as compared to HP (h,i,j). Significance was drawn by analysing the data of minimum three replicates with unpaired t-Test. (*p<0.05; **p<0.01; ***p<0.001; NS: Not significant)

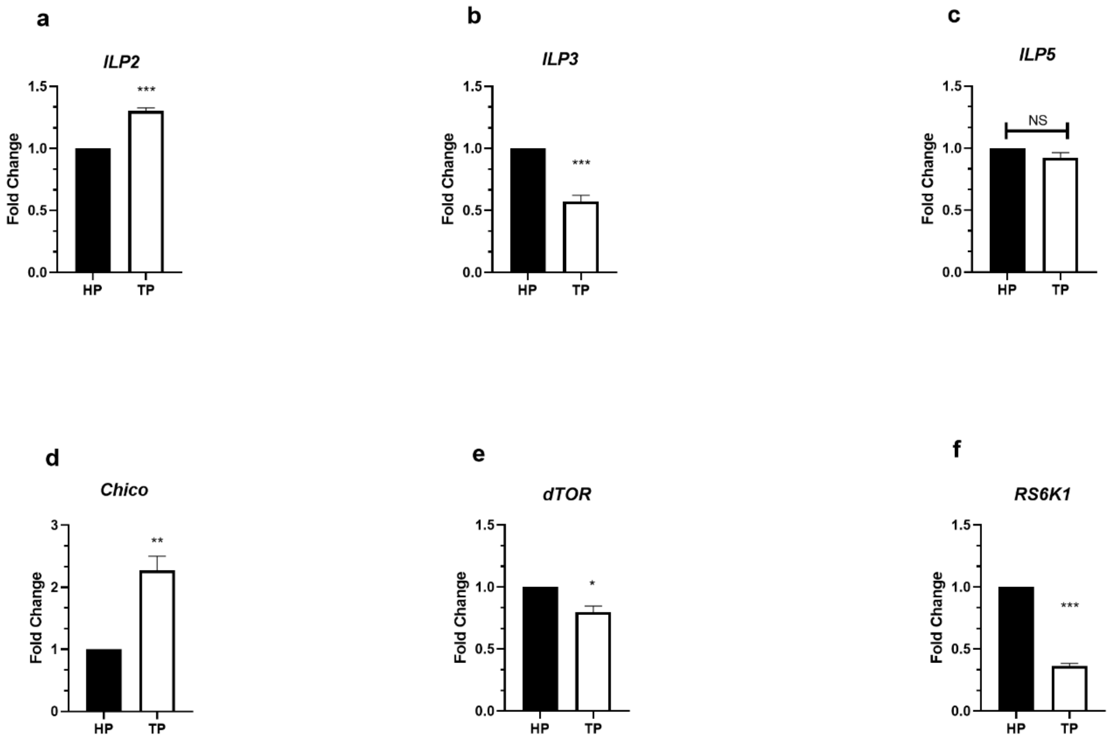


**Figure S2**: Brain-specific aging associated changes in expression of molecular players involved in *IIS-dTOR* signalling pathway. Natural aging in the brain may enhance cumulative *IIS* signalling as expression level *ILP2* was enhanced, whereas *ILP3* was downregulated and *ILP5* remained unaltered in TP brain as compared to that of HP (a,b,c). Similarly, *ILP* downstream *Chico* level was also upregulated (d). However, unlike *IIS*, the downstream *dTOR-RS6K1* signalling cascade may be repressed with natural aging as *dTOR* and RS6K1 level was inhibited in the TP brain as compared to that of HP (e,f). Significance was drawn by analysing the data of minimum three replicates with unpaired t-Test. (*p<0.05; **p<0.01; ***p<0.001; NS: Not significant)

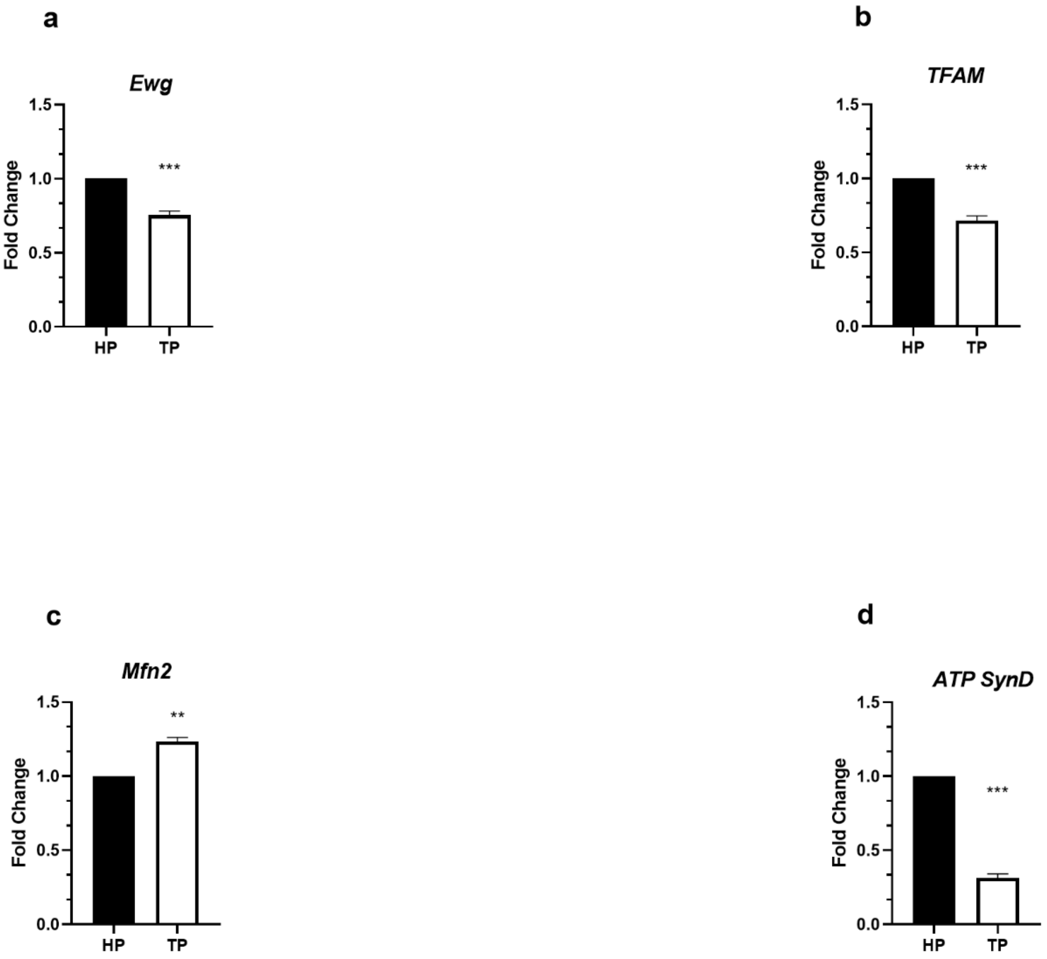


**Figure S3**: Brain-specific aging associated changes in expression of molecular players involved in mitochondrial dynamics. Natural aging in brain inhibits mito-biogenesis capacity as *Ewg* and *TFAM* expression level were inhibited in the TP brain as compared to that of HP (a,b). However, mito-quality control is enhanced with natural aging as *Mfn2* level was upregulated in the TP brain as compared to that of HP (c). Natural aging diminishes respiratory capacity in the brain as *ATP SynD* was diminished in the TP brain as compared to that of HP (d). Significance was drawn by analysing the data of minimum three replicates with unpaired t-Test. (*p<0.05; **p<0.01; ***p<0.001; NS: Not significant)

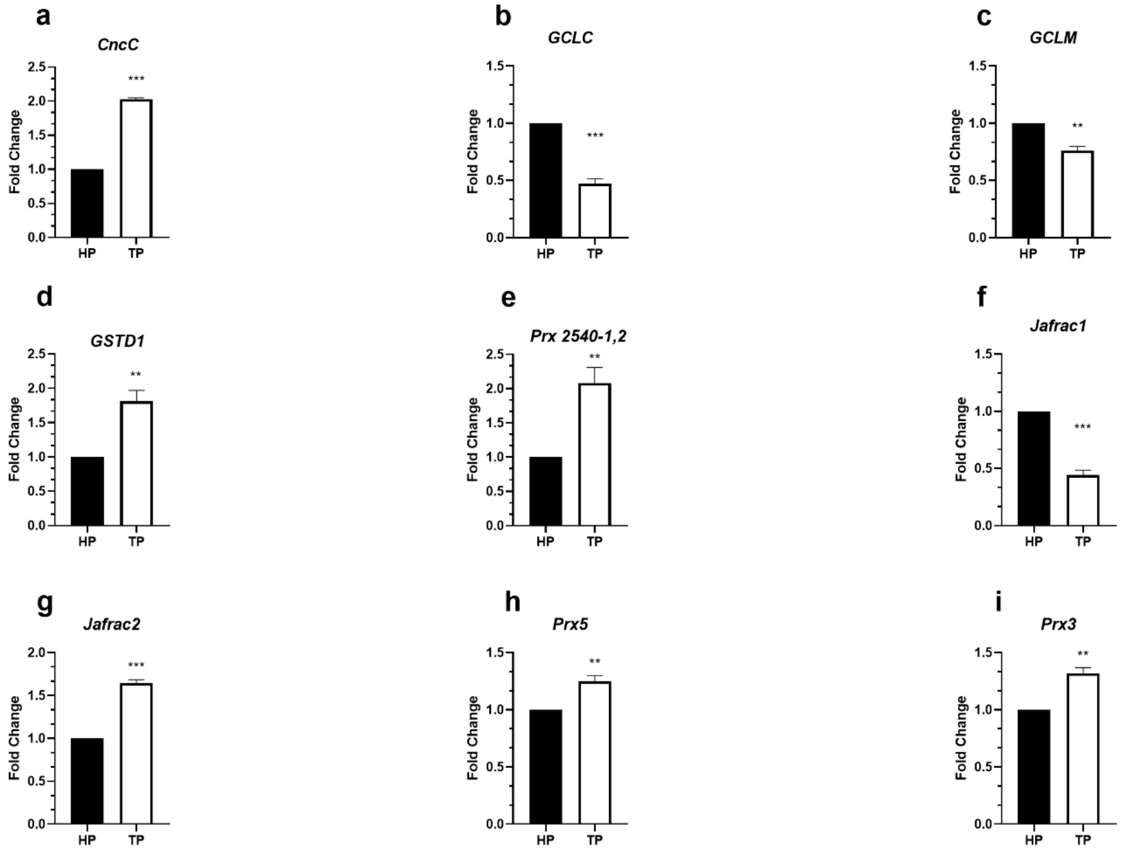


**Figure S4**: Brain-specific aging associated changes in expression of molecular players involved in phase II ADS. Natural aging in the brain upregulates phase II ADS mediator *CncC* (a). However, *CncC* downstream *GCLC* and *GCLM* were downregulated in the TP brain, suggesting a decline in GSH synthesis with aging (b,c). Further, *GSTD1*, *Prx 2540-1,2*, *Jafrac2*, *Prx5* and *Prx3* were upregulated in TP brain, whereas *Jafrac1* was downregulated (d,e,f,g,h,i). Significance was drawn by analysing the data of minimum three replicates with unpaired t-Test. (*p<0.05; **p<0.01; ***p<0.001; NS: Not significant)

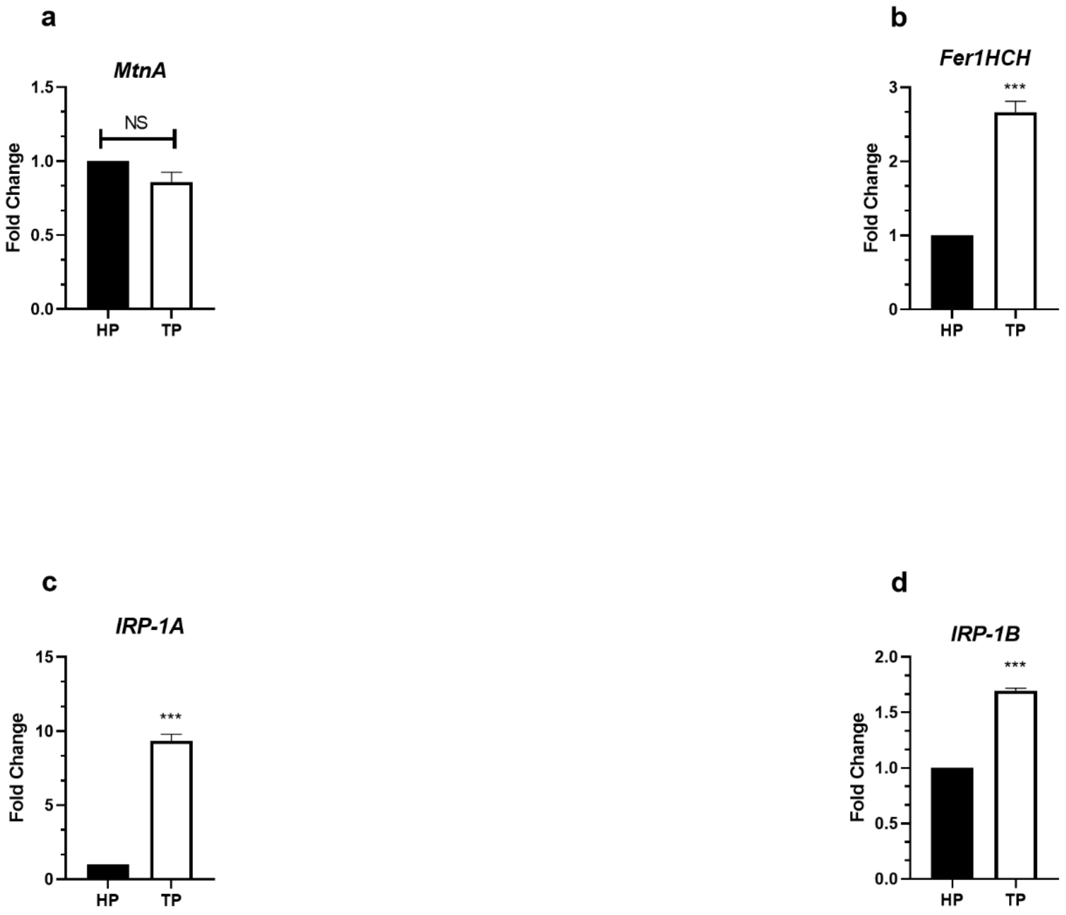


**Figure S5**: Brain-specific aging associated changes in expression of molecular players involved in metal homeostasis. Natural aging in the brain did not alter *MtnA* expression (a). But with aging iron accumulation in the brain may be enhanced and in response to that *Fer1HCH* was upregulated in TP brain (b). Similarly in the TP brain iron uptake was also enhanced as *IRP-1A* and *IRP-1B* expression was upregulated (c,d). Significance was drawn by analysing the data of minimum three replicates with unpaired t-Test. (*p<0.05; **p<0.01; ***p<0.001; NS: Not significant)
